# Methylotrophic methanogenesis in the Archaeoglobi revealed by cultivation of *Ca.* Methanoglobus hypatiae from a Yellowstone hot spring

**DOI:** 10.1101/2023.09.08.556235

**Authors:** Mackenzie M. Lynes, Zackary J. Jay, Anthony J. Kohtz, Roland Hatzenpichler

**Affiliations:** Department of Chemistry and Biochemistry, Center for Biofilm Engineering, and Thermal Biology Institute, Montana State University, Bozeman, MT 59717, USA; Department of Microbiology and Cell Biology, Montana State University, Bozeman, MT 59717, USA

**Keywords:** archaea, MCR, methane, stable isotope tracing, thermophile, transcriptomics

## Abstract

Over the past decade, environmental metagenomics and PCR-based marker gene surveys have revealed that several lineages beyond just a few well-established groups within the Euryarchaeota superphylum harbor the genetic potential for methanogenesis. One of these groups are the Archaeoglobi, a class of thermophilic euryarchaeotes that have long been considered to live non-methanogenic lifestyles. Here, we enriched *Candidatus* Methanoglobus hypatiae, a methanogen affiliated with the family Archaeoglobaceae, from a hot spring in Yellowstone National Park. The enrichment is sediment-free, grows at 64-70 °C and a pH of 7.8, and produces methane from mono-, di-, and tri-methylamine. *Ca.* M. hypatiae is represented by a 1.62 Mb metagenome-assembled genome with an estimated completeness of 100% and accounts for up to 67% of cells in the culture according to fluorescence *in situ* hybridization. Via genome-resolved metatranscriptomics and stable isotope tracing, we demonstrate that *Ca.* M. hypatiae expresses methylotrophic methanogenesis and energy-conserving pathways for reducing monomethylamine to methane. The detection of Archaeoglobi populations related to *Ca.* M. hypatiae in 36 geochemically diverse geothermal sites within Yellowstone National Park, as revealed through the examination of previously published gene amplicon datasets, implies a previously underestimated contribution to anaerobic carbon cycling in extreme ecosystems.

## Introduction

Methanogenesis is one of the most ancient metabolic pathways and plays a major role in the biogeochemical carbon cycle. Phylogenomic reconstructions and geological evidence suggest that methanogenesis was among the earliest metabolisms to evolve and that the last common ancestor of all extant archaea likely was a methanogen (1–9). Therefore, the study of methanogens is essential for understanding the co-evolution of life and the biosphere. Methanogenic archaea are the primary producers of biogenic methane (CH_4_) and contribute approximately 60% to the estimated 576 Tg of annual global methane emissions to the atmosphere (10, 11). Methanogenic pathways are classified by their carbon and electron sources (12–14). All methanogenic pathways converge at the terminal methane-forming step catalyzed by the methyl-coenzyme M reductase (MCR) complex. MCR and its homologs also catalyze the reverse reaction in the anaerobic oxidation of alkanes in alkanotrophic archaea (15, 16). MCR is uniquely present in all methanogens and is commonly used to identify potential methane and/or alkane cycling archaea in sequencing surveys (12, 17).

The physiology and biochemistry of methanogens has near-exclusively been investigated in axenic cultures of microorganisms belonging to the Euryarchaeota superphylum (12, 17–19). These predominantly grow by acetoclastic or CO_2_-reducing hydrogenotrophic methanogenesis, with only rare observations of Euryarchaeotal methyl-reducing methanogens (12, 20, 21). As a result, despite the dominance of methyl-based methanogenesis in anoxic environments with high salt and/or high sulfate concentration (*e.g.*, anoxic marine sediments, coastal wetlands, hypersaline lakes), methylotrophic methanogenesis has in the past often been considered to be of comparatively limited environmental distribution. The extensive use of environmental metagenomics has led to the discovery of metagenome-assembled genomes (MAGs) encoding MCR from new lineages that are prevalent in anoxic environments, both within and outside the Euryarchaeota (2, 12, 22–26).

The majority of MAGs affiliated with archaeal phyla outside the Euryarchaeota are predicted to be methyl-reducing methanogens, with the exception of *Candidatus* (*Ca.*) Nezhaarchaeota (25, 27) and *Ca.* Methanomixophus affiliated with the order Archaeoglobales, which have been hypothesized to be CO_2_-reducing hydrogenotrophic methanogens (12, 25, 28). This result is consistent with the observation that methylated methanogenic substrates, including methylamines and methanol, are prevalent in the environment, although their concentrations in hot springs is currently unknown. Further, methyl-reducing methanogenesis is considered the predominant mode of methanogenesis in anoxic marine, freshwater, and hypersaline sediments (reviewed in (20)).

Members of the class Archaeoglobi have long been considered non-methanogenic with isolates characterized as dissimilatory sulfate reducers brought into culture as early as 1987 (29). To date, only nine species of the class Archaeoglobi have been obtained in axenic culture, and all were sourced from marine hydrothermal systems or off-shore oil reservoirs (30). The discovery of both MCR (25, 31, 32) and methyl-H_4_M(S)PT:coenzyme M methyltransferase (MTR) complexes in genomes of the Archaeoglobaceae have suggested that members of this family may live by methanogenesis (28).

Very recently, important progress towards experimental verification of methanogenesis by members of this family has been made. Liu *et al.* reported the *in situ* expression of genes related to hydrogen-dependent methylotrophic methanogenesis and heterotrophic fermentation within populations of Archaeoglobi in a high-temperature oil reservoir (28). Lynes, Krukenberg *et al.* reported that Archaeoglobi can be enriched in hot spring mesocosms under methanogenic conditions (33). Wang *et al.* reported that *mcrABG* and other methanogenesis marker genes encoded by two Archaeoglobales MAGs were highly expressed in hot spring microcosms incubated at 65 °C and 75 °C (34). Importantly, one of these Archaeoglobales MAGs represented the only Mcr-encoding archaeon that expressed *mcrABG* genes in methanogenic microcosms performed without substrate addition or with the addition of 10 mM methanol at 75 °C. This indirectly demonstrated the methanogenic nature of this archaeon (34). Last, Buessecker *et al.* reported the establishment of a methanogenic enrichment culture of *Ca.* Methanoglobus nevadensis from Great Boiling Spring (NV, USA) (35). The culture yields up to 158 µM methane after two weeks of incubation at its optimal growth temperature of 75 °C. *Ca.* M. nevadensis is represented by a 63% complete MAG obtained from the culture and a 98% complete MAG obtained a decade earlier (35).

Here, we report on the enrichment cultivation of *Ca.* Methanoglobus hypatiae LCB24, a methanogen affiliated with the family Archaeoglobaceae, from a hot spring in Yellowstone National Park (YNP). Using a combination of targeted cultivation, growth experiments, microscopy, stable isotope tracing, metagenomics, and metatranscriptomics, we demonstrate that *Ca.* M. hypatiae lives by methylotrophic methanogenesis and converts different methylamines to methane. By examining previously published datasets for the presence of Mcr-encoding Archaeoglobi, we demonstrate that these archaea are distributed in geothermal features of YNP, where they likely contribute to anaerobic carbon cycling. Our study presents direct evidence of methanogenesis within the Archaeoglobaceae and adds to the growing body of evidence demonstrating that methanogenesis is widely spread within the Euryarchaeota superphylum.

## Materials and Methods

All chemicals used in this study were sourced from Sigma Aldrich unless otherwise specified.

### Sample Collection, Enrichment, and Cultivation

In November 2021, a slurry of sediment and water (1:9) was collected from an unnamed hot spring in the Lower Culex Basin of Yellowstone National Park (YNP), WY, USA. In our previous survey of Mcr-encoding archaea in YNP (33), this hot spring was given the identifier LCB024 (44.573294, −110.795388; pH 7.8, 67 °C). A mixture of surface sediment (∼1-2 cm deep) and hot spring water was collected into a glass bottle and sealed headspace-free with a thick butyl rubber stopper. Collected material was transported back to the lab and stored at room temperature. Using this material as inoculum, 30 mL enrichments were established in February 2022 in 60 mL serum bottles. Material was homogenized by mixing and was then diluted 1:10 (v/v) with anoxic medium in an anoxic glove box (N_2_/CO_2_/H_2_; 90/5/5%).

Medium was prepared anoxically as described previously (36). Basal mineral medium contained a base of (per liter): KH_2_PO_4_, 0.5 g; MgSO_4_·7H_2_O, 0.4 g; NaCl, 0.5 g; NH_4_Cl, 0.4 g; CaCl_2_·2H_2_O, 0.05 g; HEPES, 2.38 g; yeast extract, 0.1 g; and 0.002% (w/v) (NH_4_)_2_Fe(SO_4_)_2_·6H_2_O. Medium was transferred to a Duran flask with a side opening and autoclaved for 20 m at 121 °C. Medium was then further supplemented with 5 mM NaHCO_3_, 1 mL trace element solution SL-10, 1 mL Selenite-Tungstate solution, 1 mL CCM vitamins (37), 0.0005% (w/v) resazurin, 10 mg of coenzyme-M, 2 mg sodium dithionite, 1 mM dithiothreitol, 1 mM Na_2_S·9H_2_O, with pH adjusted to 7.8 using sodium hydroxide (NaOH, 12 N). Serum bottles were sealed with butyl rubber stoppers and aluminum crimps before the headspace was exchanged with N_2_ (99.999%) for 5 minutes and set to a 200 kPa N_2_ atmosphere. Monomethylamine (MMA) was added from a filter-sterilized methylamine-hydrochloride anoxic stock solution to a final concentration of 10 mM. The bacterial antibiotics streptomycin (50 mg/L; inhibitor of protein synthesis) and vancomycin (50 mg/L; inhibitor of peptidoglycan synthesis) were added from filter-sterilized anoxic stock solutions. The enrichments were incubated at 70 °C in the dark without shaking. Cultures were maintained by regular transfer of 10% v/v into fresh media, which contained MMA and antibiotics. A sediment-free culture was obtained after the third transfer after which it was transferred at 10% v/v to 50 mL in 125 mL serum bottles.

### Stable Isotope Tracing

The conversion of ^13^C- or D_3_-MMA (^13^CH_3_-NH_2_, CD_3_-NH_2_) to ^13^CH_4_ or CD_3_H was tracked by incubating active enrichment cultures in the presence of 20% labeled substrate (98%; Cambridge Isotope Laboratories). Incubations were carried out in 30 mL culture volumes in 60 mL serum bottles with 8% v/v inoculum, 50 mg/L streptomycin, 50 mg/L vancomycin, 10 mM MMA, and N_2_ gas (99.999%) incubated in anoxic media (pH 7.8, 70 °C) in six replicates (SI Appendix, Fig. S3). Duplicate control incubations included (i) ^12^C-MMA and (ii) inoculum without MMA. Triplicate control incubations were performed with (iii) ^12^C-MMA plus 10 mM bromoethanesulfonate (BES) added in mid-exponential phase (day 33) to inhibit methanogenesis and (iv) 10 mM BES added at time of inoculation (day 0) without substrate. Headspace samples were collected throughout the experiment as described above and analyzed using a Shimadzu QP2020 NX GCMS equipped with a GS-CarbonPLOT column (30 m × 0.35 mm; 1.5 μm film thickness; Agilent) and operated in Selected Ion Monitoring mode. The instrument was operated using the method described in Ai et al., 2013 (38) with helium as a carrier gas. All injections were performed by a Shimadzu AOC-6000 autosampler robot. Peak areas corresponding to m/z ratios of 16 for ^12^CH_4_, 17 for ^13^CH_4_, and 19 for CD_3_H were used for quantification.

### Metagenomic Sequencing, Assembly, and Annotation

Two metagenomes were obtained over the course of this study. A 42 mL aliquot of the fourth transfer of the enrichment (Fig. 1 T4-MG) was filtered onto a 0.22 µm filter. The filter was transferred to a lysing matrix E tube and DNA extracted immediately following filtration. Genomic DNA was extracted using the FastDNA Spin Kit for Soil (MP Biomedicals, Irvine, CA) following the manufacturer’s guidelines.

**Fig. 1.**
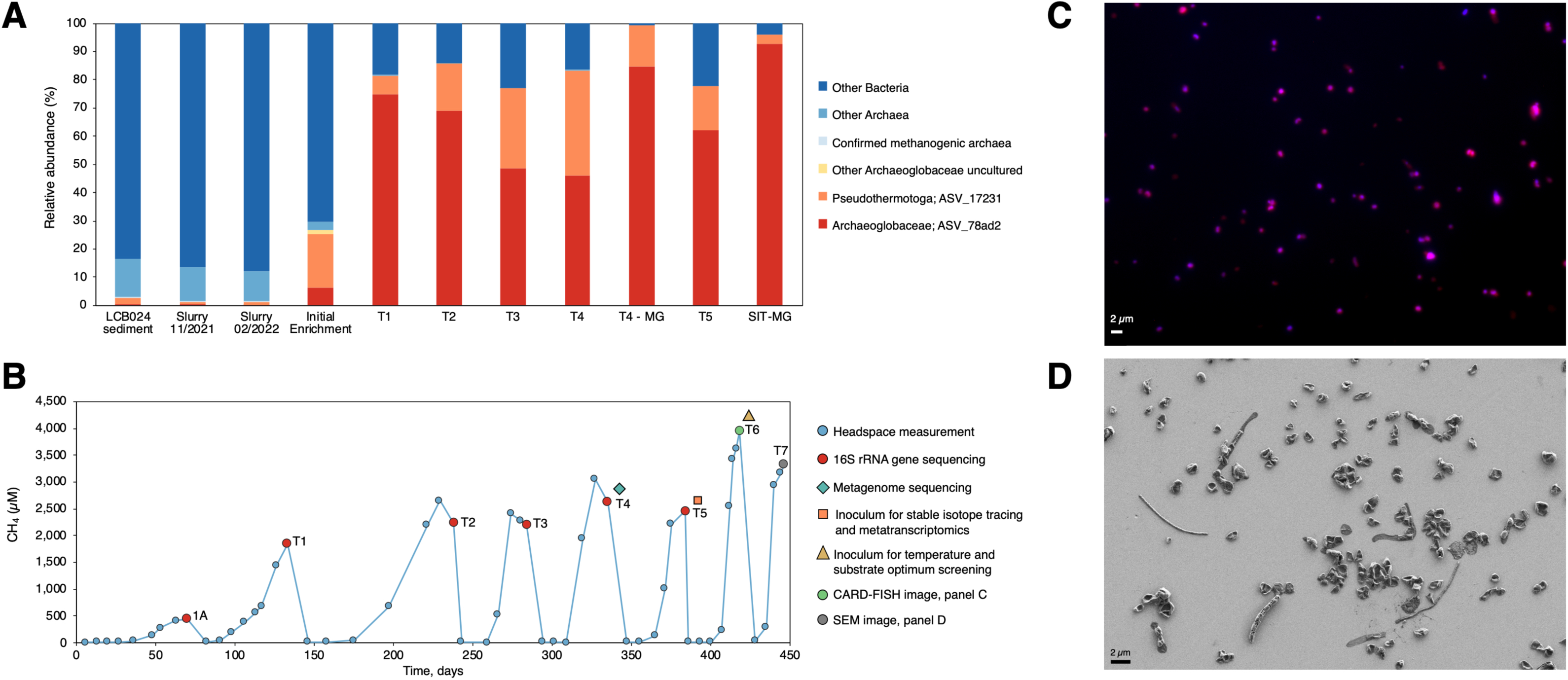
Community composition and methane production of the methanogenic enrichment culture containing *Ca.* Methanoglobus hypatiae LCB24. *(A)* Relative abundance of 16S rRNA gene amplicons in the initial sediment from hot spring LCB024, the slurry collected in November 2021, slurry material used to initiate enrichments in February 2022, the initial enrichment, and five subsequent transfers (T1-T5) are shown. For comparison, the estimated relative abundance of two metagenomic samples (T4-MG and SIT-MG) is included. The metagenome recovered from a replicate from the stable isotope tracing experiment incubated in the presence of deuterated methylamine (SIT-MG) revealed *Ca.* M. hypatiae grew to 92.8% relative abundance during the experiment. The two most abundant ASVs across enrichment transfers are shown with other taxa collapsed. Other methanogenic archaea were not identified in the initial enrichment or in any subsequent transfers. Relative sequence abundance for all ASVs is reported in SI Appendix, Table S1. *(B)* Headspace methane produced over long-term cultivation. The time between transfers decreased while the average maximum concentration of methane increased over time. Culture 1A represents the initial enrichment. A history of methane measurements can be found in SI Appendix, Table S2. *(C)* Visualization of *Ca.* M. hypatiae cells at T6 labeled via CARD-FISH by the general archaea probe Arch915 (red). DAPI staining of cells is in blue. *(D)* Cell morphologies in enrichment culture LCB24 at T7 as observed by scanning electron microscopy.

A second metagenome was recovered from one of the six culture replicates grown in the presence of CD_3_-NH_2_ and used for recruiting transcriptomic reads from the other replicates (Fig. 1 SIT-MG). A 60 mL syringe flushed with N_2_ gas was used to transfer 30 mL of culture to a sterilized oak ridge tube. Cells were harvested through centrifugation for 30 minutes at 10,000 rpm at 4 °C. The supernatant was removed, and DNA extracted from the pellet using the FastDNA Spin Kit for Soil (MP Biomedicals, Irvine, CA) following the manufacturer’s guidelines. Genomic DNA for both metagenomes was shipped to SeqCenter (Pittsburgh, PA) and sample libraries were prepared using the Illumina DNA Prep kit and 10 bp unique dual indices (UDI). The first metagenome (T4-MG) was sequenced on a NextSeq 2000 System (Illumina) and the second (SIT-MG) sequenced on a NovaSeq 6000 System (Illumina), each producing 2×151 bp reads. Demultiplexing, quality control, and adapter trimming was performed with bcl-convert v3.9.3. Quality of the reads were evaluated with FastQC before quality, linker and adapter trimming, artifact and common contaminant removal, and error correction were performed with the rqcfilter2 pipeline (maxn=3, maq=10, trimq=20) and bbcms (mincount=2, hcf=0.6). Resulting reads were assembled with SPAdes v3.15.13 (Nurk, 2017) (-k 33,55,77,99,127 --meta –only-assembler) and coverage was determined with bbmap v38.94 (ambiguous=random) (https://sourceforge.net/projects/bbmap) (39). In addition to the initial assembly, co-assemblies using both T4-MG and SIT-MG metagenomes were also performed (1) with reads directly fed into SPAdes with the –only-assembler option excluded; and (2) with the trimmed and error corrected reads and the same SPAdes parameters as above. The statistics of MAGs generated through various assembly and quality control methods were evaluated, and the approach that produced the highest quality MAG was chosen for subsequent analysis (Dataset S1). Quality was determined by considering factors such as the number of resulting sequences, total length, completeness, and the minimum, maximum, and average sequence lengths. Annotation of the assembled sequences was performed with Prokka v1.14.16 (40). Assembled scaffolds ≥2000 bp were binned using Maxbin v2.2.7 (41), Metabat v2.12.1 (with and without coverage) (42), Concoct v1.0.0 (43), Autometa v1 (bacterial and archaeal modes with the machine learning step) (44), followed by bin refinement with DAS_Tool v1.1.6 (45), as previously described (46). Bin completeness and redundancy were assessed with CheckM v1.2.2 (47).

### RNA Extraction, Sequencing, and Transcriptomic Processing

Total RNA was extracted for transcriptomics from four of the six replicates of Archaeoglobus cultivated in the presence of labeled substrate (^13^CH_3_-NH_2_ or CD_3_-NH_2_) for a total of eight replicates. Each replicate culture in the exponential growth phase (day 32) was moved from the 70 °C incubator to an ice bath placed at −20 °C for 40 minutes to stop cellular activity. A 60 mL syringe flushed with N_2_ gas was used to transfer 30 mL of culture to a sterilized oak ridge tube and kept on ice. Cells were harvested through centrifugation for 30 minutes at 10,000 rpm at 4 °C. The supernatant was removed, and the pellet transferred to a lysing matrix E tube (MP Biomedicals, Irvine, CA) to which 600 µL of RNA lysis buffer was added. Samples were homogenized in a MP Bioscience FastPrep instrument for 40 seconds at a speed setting of 6.0 m/s followed by centrifugation for 15 minutes at 14,000 rpm. RNA was extracted using the Quick-RNA miniprep kit (Zymo Research, Irvine, CA) including a DNAse treatment step and eluted in 50 µL of RNAse free water. Centrifugation steps were performed at 15,000 rpm and the final spin for elution at 10,000 rpm. Of the eight replicates extracted, six measured >50 ng/µL (3x ^13^CH_3_-NH_2_ and 3x CD_3_-NH_2_) and were sent for transcriptomic sequencing at SeqCenter (Pittsburgh, PA). Samples were DNAse treated with Invitrogen DNAse (RNAse free). Library preparation was performed using Illumina’s Stranded Total RNA Prep Ligation with Ribo-Zero Plus kit and 10bp UDI. Sequencing was done on a NovaSeq 6000, producing paired end 151bp reads. Demultiplexing, quality control, and adapter trimming was performed with bcl-convert (v4.1.5). Read quality was further evaluated with FastQC v0.11.9 (48) before quality trimming and artifact, rRNA, and common contaminant removal with the rqcfilter2 pipeline (trimq=28, maxns=3, maq=20), and error correction with bbcms (mincount=2, hcf=0.6) from the BBTools suite v38.94 (39). Additional rRNA gene reads were detected and removed with Ribodetector v0.2.7 (49) and any remaining rRNA gene reads were finally removed with bbmap, using rRNA genes recovered from the metagenomes (see below) as references. The resulting mRNA reads were mapped against annotated genes from the paired metagenomes with bbmap to calculate RPKM (ambig=random).

### Data Availability

All metagenomic, metatranscriptomic, and amplicon data discussed in this manuscript are available under NCBI BioProject ID PRJNA1014417. McrA gene amplicon data from YNP hot springs discussed in this manuscript has been previously published (Lynes et al., 2023) and is available under NCBI under BioProject PRJNA859922.

## Results and Discussion

### Cultivation

In our recent survey on the diversity of Mcr-encoding archaea in the geothermal features of YNP, mesocosms seeded with biomass from a hot spring located within the Lower Culex Basin (LCB024; pH 7-8, 56-74 °C), had shown potential to enrich for methanogenic Archaeoglobi (33). Using a sediment slurry collected from LCB024, we initiated incubations supplied with monomethylamine (MMA) and antibiotics incubated in anoxic media (pH 7.8, 70 °C) under a N_2_ headspace. The relative abundance, as determined by 16S rRNA gene amplicon sequencing, of Archaeoglobi-affiliated organisms in LCB024 was 0.32% in the initial slurry and had fallen to 0.02% by the time incubations were initiated a few months after samples had been collected (Fig. 1A).

Methane was detected after 36 days in the initial enrichment and the culture transferred to fresh media after reaching the late exponential phase of methane production following 70 days of incubation (447 µM; Fig. 1B). Five Archaeoglobi related 16S rRNA gene amplicon ASVs were identified in the initial enrichment, however one ASV grew to dominate the microbial community after the first transfer and reached 74.8% relative abundance after 62 days. In the transfers that followed, Archaeoglobi-related sequences became the only archaeal reads detected by 16S rRNA gene amplicon sequencing with the second most abundant organism a bacterium affiliated with the *Pseudothermotoga* at 6.8%. Although the CO_2_-reducing methanogen *Methanothermobacter* sp. was detected at 0.45% relative abundance in the slurry material used for inoculation, it was not detected in any subsequent transfers, nor were any known methanogens. Over subsequent transfers (238 days, T2-T5), the relative abundance of Archaeoglobi ASVs ranged from 46 to 69% and the final methane yield steadily increased from 1,844 to 2,459 µM. A sediment-free enrichment was obtained by the third transfer. Starting with the fourth transfer, the culture volume was scaled from 30 mL to 50 mL. By the sixth transfer, the culture produced 3,943 µM methane within 34 days. Metagenomic sequencing at two timepoints (day 335 of the enrichment and day 33 of the isotope tracing experiment described below) and 16S rRNA gene amplicon sequencing over recurring transfers (Fig. 1A) demonstrated that ASVs and MAGs affiliated with Archaeoglobi represented the only archaeon in culture LCB24. A single MCR complex (*mcrAGCDB*) belonging to the Archaeoglobi MAG was present, indicating this MAG represents the only methanogenic population.

### Metagenomics and Phylogenetics

The reconstructed Mcr-encoding Archaeoglobi MAG from culture LCB24 was 1.62 Mbp in length with an estimated completeness of 100% according to checkM (SI Appendix, Table S3). This MAG was the result of a combined assembly of the T4-MG and SIT-MG metagenomes as this method yielded an improved assembly. Therefore, it was used for phylogenomic analysis against Archaeoglobi reference MAGs and genomes using 33 conserved single copy marker proteins and 16S rRNA genes (Fig. 2AB, SI Appendix, Table S4). The phylogenomic analysis showed that MAGs encoding MCR complexes clustered separately from those lacking *mcr* gene sequences. Consistently, 16S rRNA gene phylogeny supported this clustering with a pronounced separation of hot spring reference genomes and MAGs from current known isolates of Archaeoglobi, resulting in three main clusters: (i) those retrieved from North American hot springs (YNP and Great Boiling Spring, GBS), (ii) those originating from hot springs in China, and (iii) isolates, all of which were obtained from deep-sea marine hydrothermal systems (Fig. 2B).

**Fig. 2.**
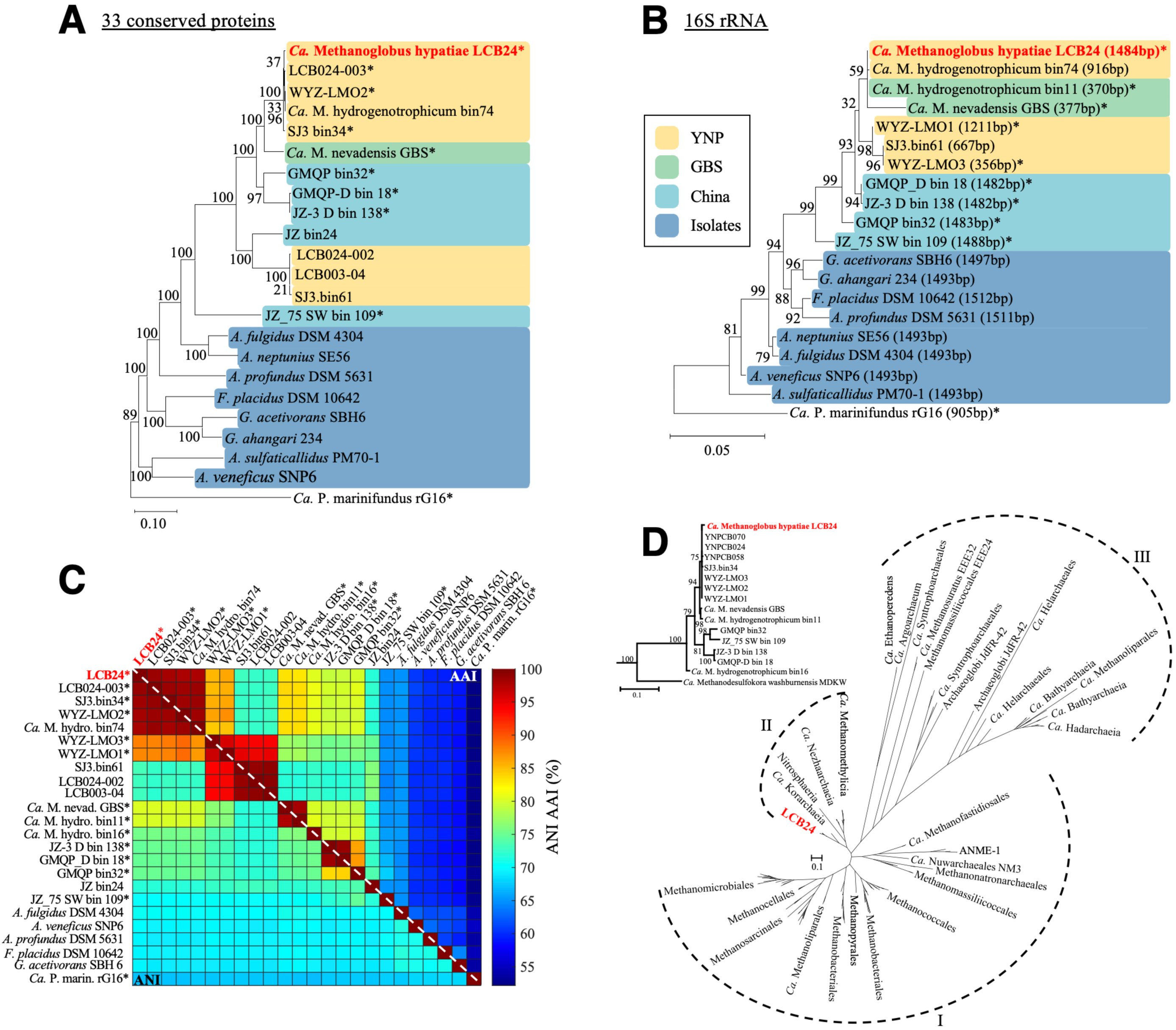
Phylogenetic affiliation of *Ca.* M. hypatiae LCB24. *(A)* Maximum-likelihood tree, inferred with fasttree and WAG model (midpoint root), using a concatenated alignment of 33 conserved single copy proteins (list provided in SI Appendix, Table S4). References are colored by the habitat or type from which sequences had been recovered: hot springs in Yellowstone National Park (YNP), yellow; Great Boiling Spring (GBS), green; hot springs in China, blue; isolates from marine hydrothermal vent systems, dark blue. *(B)* Maximum-likelihood tree inferred with fasttree using 16S rRNA genes with length in base pairs (bp). *(C)* ANI and AAI analysis of reference Archaeoglobales MAGs and genomes. Asterisks (*) indicate MAGs containing *mcrA*, apart from the MAG of *Ca.* M. hydrogenotrophicum bin74 which encodes a *mcrA* that is interrupted by a stop codon. AAI and ANI values are provided in SI Appendix, Fig. S1. *(D)* Maximum-likelihood tree, inferred with IQtree2 and the LG+C60+F+G model, from the amino acid alignment of McrA. Dashed lines indicate McrA/AcrA groups: (I) McrA from methanogens and ANME (MCR-type), (II) McrA from TACK lineages (MCR-type), (III) McrA-like from proposed and experimentally confirmed alkane oxidizing archaea (ACR-type). Insert shows MAGs closely related to *Ca.* M. hypatiae LCB24.

LCB24 and closely related reference MAGs and isolate genomes exhibited a range of amino acid identities (AAI, 52.6-98.6%; Fig. 2C). Altogether, the LCB24 MAG was found to be highly related to previously obtained Archaeoglobi MAGs encoding the MCR complex and only distantly related to other Archaeoglobales sp. confirmed sulfate-reducing Archaeoglobales (AAI, 58.9-65%; ANI, 70.3-70.6%; 16S rRNA ANI, 91.6-93.8%; SI Appendix Fig. S1, S2). Based on AAI, MAG LCB24 was most closely related to Archaeoglobi LCB024-003 MAG (AAI, 98.6%), which we had obtained from the same hot spring in a previous study (33). The ANI and AAI values to the closest cultured methanogen, *Ca.* Methanoglobus nevadensis GBS, are 80.2 and 83.3%, respectively. Based on these results, we designate this archaeon *Ca.* Methanoglobus hypatiae strain LCB24, named after the philosopher Hypatia of Alexandria (for a protologue, see the SI Appendix, Results and Discussion). The estimated relative abundance of *Ca.* M. hypatiae based on the SIT-MG was 92.8%. Other community members in the LCB24 culture with >1% relative sequence abundance included members of the *Pseudothermotoga* (3.2%), *Desulfovirgula* (1.7%), and the family Moorellaceae (1.3%) (Fig. 1A, Dataset S1).

The only *mcrAGCDB* genes recovered from both metagenomes belong to the genome of *Ca.* M. hypatiae. Phylogenetic analysis of the single copy of McrA indicated its close relationship to McrA sequences found in members of the TACK superphylum (Fig. 2D). This contrasts with the placement of *Ca.* M. hypatiae within the Euryarchaeota based on phylogenomics (Fig. 2A), suggesting that Archaeoglobi could have obtained the MCR complex as a result of a horizontal gene transfer event from an archaeon in the TACK superphylum (7, 8). Also, it could indicate that non-methanogenic Archaeoglobi lost the capacity for anaerobic methane cycling after they had diverged from a shared methanogenic ancestor.

### Methanogenic Activity of *Ca.* M. hypatiae

To gain insight into the activity of *Ca.* M. hypatiae under methanogenic and non-methanogenic conditions, a stable isotope tracing (SIT) experiment was conducted. Cultures were incubated in the presence of 10 mM of MMA; 8 mM of substrate were isotopically light, whereas the remaining 2 mM consisted of either ^13^C-monomethylamine (^13^CH_3_-NH_2_) or D_3_-monomethylamine (CD_3_-NH_2_). Addition of the methanogenesis inhibitor bromoethanesulfonate (BES) was used as a non-methanogenic control (Fig. 1B, 3, SI Appendix, Fig. S3). On average across six replicates, the cultured converted ^13^CH_3_-NH_2_ to 356 µM ^13^CH_4_ (17.8%) and 138.71 µM ^13^CO_2_ (6.9%) by day 32 (Fig. 3AC, Dataset S2). The conversion of CD_3_-NH_2_ was nearly identical yielding 355 µM CD_3_H (Fig. 3B). In the exponential phase of methane production, five of the six replicates were harvested for metagenomic and metatranscriptomic sequencing while the sixth replicate was allowed to grow to stationary phase. The replicate allowed to grow in each respective experiment converted the provided ^13^CH_3_-NH_2_ to 717.7 µM ^13^CH_4_ (35.9%) and 212.95 µM ^13^CO_2_ (10.65%) or CD_3_-NH_2_ to 394.76 µM CD_3_H (19.7%) by day 38 (Fig. 3ABC). These results confirmed monomethylamine was converted to methane by the LCB24 culture. The production of ^13^CO_2_ may represent the dismutation of ^13^CH_3_-NH_2_ to generate reducing power for methanogenesis via the methyl-branch of the Wood-Ljungdahl pathway or may be explained by other organisms in the culture catabolizing MMA. Yet, no transcriptomic evidence for this activity was present in this experiment. No methane production was observed for cultures treated with BES or in cultures incubated without MMA (Fig. 3D). When BES was added to cultures in the exponential phase, methane production ceased indicating the generation of methane is reliant on the Archaeoglobi MCR (Fig. 3E).

**Fig. 3.**
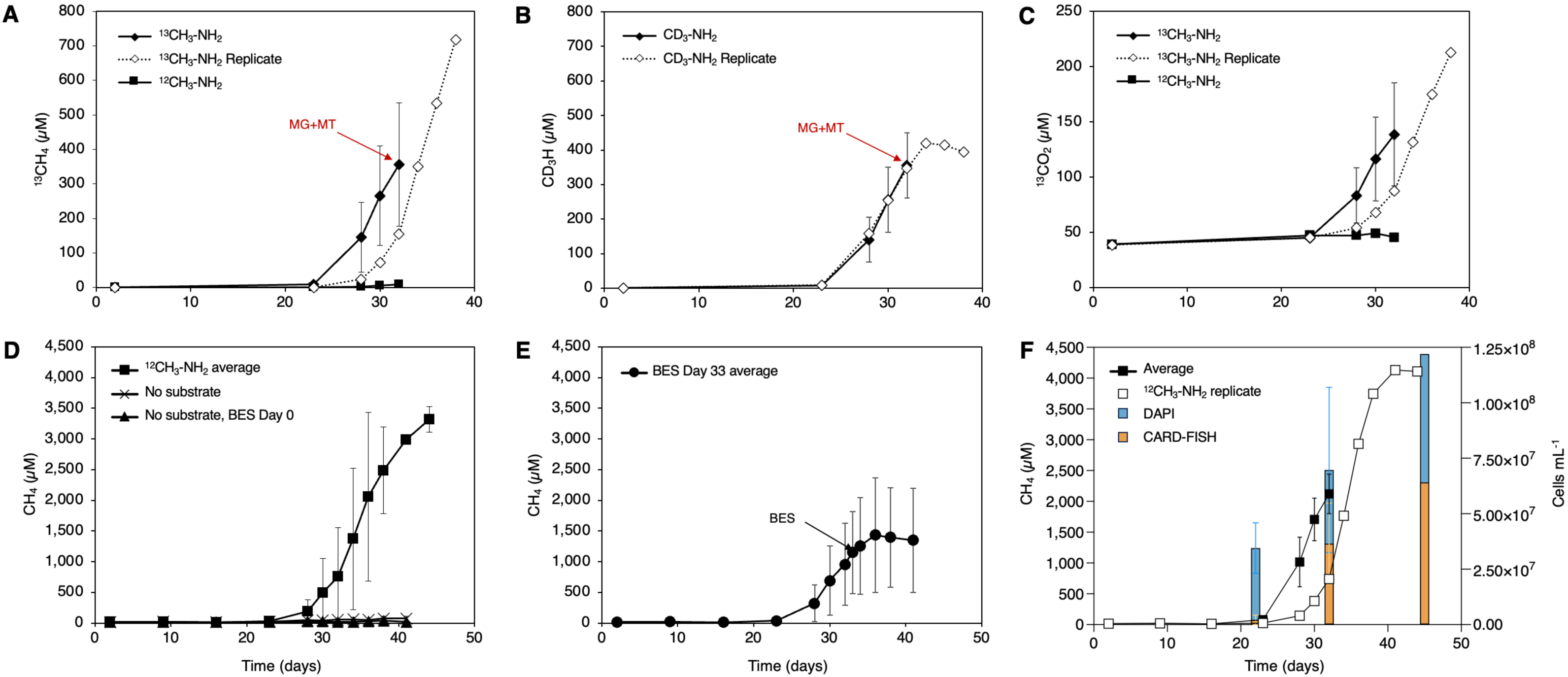
Conversion of stable isotope labeled monomethylamine to methane by culture LCB24. *(A)* Production of ^13^CH_4_ in cultures amended with ^13^CH_3_-NH_2_ vs. ^12^CH_3_-NH_2_ (6 replicates). *(B)* Production of CD_3_H in cultures amended with CD_3_-NH_2_ (6 replicates). *(C)* Production of ^13^CO_2_ in cultures amended with ^13^CH_3_-NH_2_ vs. ^12^CH_3_-NH_2_. For plots A and B, ten total replicates across treatments were sacrificed during mid-exponential phase for metagenomic or metatranscriptomic sequencing indicated by red arrows. ^13^CH_4_, CD_3_H, or ^13^CO_2_ production for the replicate allowed to reach stationary phase is shown as a dashed line through open diamond symbols. *(D)* Production of ^12^CH_4_ in cultures amended with ^12^C-monomethylamine (^12^CH_3_-NH_2_; 2 replicates). Cultures incubated without substrates (2 replicates) and those to which the inhibitor BES was added on day 0 (3 replicates) did not produce ^12^CH_4_ over the course of the experiment. *(E)* Production of ^12^CH_4_ in cultures amended with ^12^CH_3_-NH_2_ to which BES was added on day 33 of incubation (black arrow; 3 replicates). The average production of ^12^CH_4_ leveled off and ceased after the introduction of BES, indicating methane generation by *Ca.* M. hypatiae is MCR-dependent. Error bars indicate standard deviation of biological replicates when applicable. Measurements of ^12^CH_4_, ^13^CH_4_, CD_3_H, and ^13^CO_2_ for all replicates and controls are reported in Dataset S2. ^12^CH_4_ measurements for all controls and replicates are shown in SI Appendix, Fig. S4 and Dataset S3. *(F)* ^12^CH_4_ production and fraction of *Ca.* M. hypatiae cells in biological replicates incubated with ^13^CH_3_-NH_2_. Relative abundance of cells was determined at three time points (day 22, 32, 45) based on the fraction of *Ca.* M. hypatiae specific CARD-FISH counts (orange) versus total counts of DAPI stained cells (blue). Error bars indicate the standard deviation for four biological replicates on days 22 and 32.

### Visualization and Cell Enumeration

The growth of *Ca.* M. hypatiae was tracked in four replicates during the SIT experiment with catalyzed reporter deposition fluorescence *in situ* hybridization (CARD-FISH) using a general archaea-targeted probe Arch915 (50) and DNA-staining (DAPI) (Fig. 1C). As the production of methane increased throughout the experiment, there was a concurrent rise in the relative cell abundance of *Ca.* M. hypatiae (Fig. 3F, SI Appendix, Table S5). The initial assessment on day 22 across four replicates revealed the total cell density to be 3.45 × 10^7^ ± 1.14 × 10^7^ before substantial concentrations of methane had been detected in the headspace (<132 µM). By day 32, methane concentrations reached 1,777±739 µM and the total cell density increased to 6.97 × 10^7^ ± 3.73 × 10^7^ cells mL^-1^ with 54% (±9.6%) of cells labeled as *Ca.* M. hypatiae (Fig. 3F). All but one of these replicates were then sacrificed for further analysis. Finally on day 45, the remaining replicate reached a headspace methane concentration of 4,109 µM and a total cell density of 1.22 × 10^8^ with 53% of cells labeled as *Ca.* M. hypatiae.

Visualization of the enrichment culture via scanning electron microscopy (SEM) revealed that most cells exhibited a regular to irregular coccoid morphology, with a width ranging from 0.5-1 µm (Fig. 1D). This morphology has previously been described for other Archaeoglobi species (30, 51–53).

### Alternative Substrates and Temperature Optimum

We determined the substrate and temperature range of *Ca.* M. hypatiae by growing the culture in the presence of several substrates at 70 °C or with 10 mM MMA at 60-85 °C (Fig. 4AB). Conditions that lead to the production of methane included 10 mM trimethylamine (TMA), 10 mM dimethylamine (DMA), 10 mM MMA in media without yeast extract, and the control with 10 mM MMA and 0.01% yeast extract.

Methane production of cultures grown with MMA in the presence or absence of yeast extract were indistinguishable (5,202±606 and 5,703±410 µM CH_4_, respectively) indicating that yeast extract is not essential for methanogenic growth. Observed methane concentrations were higher in incubations amended with DMA (10,115±836 µM CH_4_) and TMA (9,524±3,626 µM CH_4_, with a wide range of 5,361-11,993 µM) on average more than the MMA controls, consistent with what has been observed for other methylotrophic methanogens (54). Incubations amended with 10 mM methanol (MeOH) did not produce methane after 47 days of incubation at 70 °C. Due to its use by sulfate reducing organisms as an electron donor (55), 10 mM lactate (LAC) was tested, as well as 10 mM MMA with 10 mM LAC, but none of these incubations produced methane. Production of methane has not been observed in any attempted transfers where hydrogen (99.9999% purity) was present in the headspace, or hydrogen with MMA was added.

The enrichment grew optimally at both 64 and 70 °C with relative amounts of methane produced at 5,304±451 µM and 5,202±606 µM, respectively. This deviates from the predicted optimal growth temperature of 74.4°C, which was derived from the translation of proteins in the Ca. M. hypatiae MAG using Tome (56). This is lower than the observed range of growth and optimum temperatures for type strains of non-methanogenic Archaeoglobus which have been demonstrated to grow between 50 and 95 °C with optimal temperatures between 75-83 °C in organisms sourced predominantly from deep sea vent environments (30). No methane production was detected at temperatures 77 °C or above or lower than 64 °C after 47 days of incubation (Dataset S4).

### Genomic and Transcriptomic Basis for Methanogenesis

The assembled metagenome obtained at the end of the SIT experiment was used to align a total of 23,376,154 metatranscriptome mRNA reads obtained from six replicates harvested in the exponential growth phase and to create a detailed reconstruction of the metabolism of *Ca.* M. hypatiae (Fig. 3AB, 5, Dataset S5). A total of 22,891,651 reads, *i.e.,* 97.8% of all recovered reads, were recruited to the *Ca.* M. hypatiae MAG. Only 2.1% of the total mRNA reads (484,503) were aligned with other co-enriched organisms. Among these, only 13 genes across four MAGs were expressed above 200 reads per kilobase of transcript per million mapped reads (RPKM) and just five genes exceeded >1,000 RPKM. Genes required for the conversion of methylamine to methane were among the top 2% of highest expressed genes transcribed by *Ca.* M. hypatiae, including genes encoding the MCR complex (*mcrAGCDB*; 13,046-18,098 RPKM), one of three monomethylamine methyltransferase copies (*mtmB*; 9,884 RPKM), dimethylamine corrinoid (*mtbC*; 3,677 RPKM), and methanol:coenzyme M methyltransferase (*mtaA*; 12,577 RPKM) (Fig. 5). Seven copies of substrate-specific methyltransferases for MMA (*mtmB*; 3 copies), DMA (*mtbB*; 2 copies), and TMA (*mttB*; 2 copies) were present in the genome, but methanol methyltransferase (*mtaB*) was not identified. These genes were differentially expressed with one copy for each type of methylamine expressed above 3,200 RPKM. In addition to *mtbC*, two gene copies of the trimethylamine corrinoid protein (*mttC*) were found in the genome but their expression was relatively low (<460 RPKM average). Monomethylamine corrinoid (*mtmC*) or methanol corrinoid (*mtaC*) proteins were not identified in *Ca.* M. hypatiae. Additionally, genes were expressed for pyrrolysine synthesis (*pylBCD*; 819, 343, 37 RPKM) and the methyltransferase corrinoid activation protein (*ramA*; 1,076 RPKM), both of which support methylamine methyltransferases in methylotrophic methanogenesis (57, 58). The absence of *mtmC* and the high expression levels of *mtbC* (3,677 RPKM) and *mtaA* (12,577 RPKM) suggest that they are responsible for the transfer of a methyl group from monomethylamine to coenzyme M (CoM) after it has been transferred by a substrate-specific methyltransferase (*mtmB*). Consistent with the observed methane production from DMA and TMA, *Ca.* M. hypatiae can use these methylamines and expressed the corresponding genes (*mtbB*, *mttB*) at comparatively high levels (JOOIALLP_01813 *mtbB* 3,249 RPKM; JOOIALLP_01787 *mttB* 5,324 RPKM; Fig. 4B, 5). It is worth noting that the expression of mtbB/mttB was detected despite the culture not having been previously exposed to DMA or TMA at the time of the transcriptomics experiment. We hypothesize that *Ca.* M. hypatiae could employ one of two strategies: it either (i) constitutively expresses all substrate-specific methyltransferases and corrinoid proteins as a precautionary measure to accommodate substrates potentially encountered *in situ*, or (ii) *Ca.* M. hypatiae transcriptionally co-regulates the genes responsible for these functions.

**Fig. 4.**
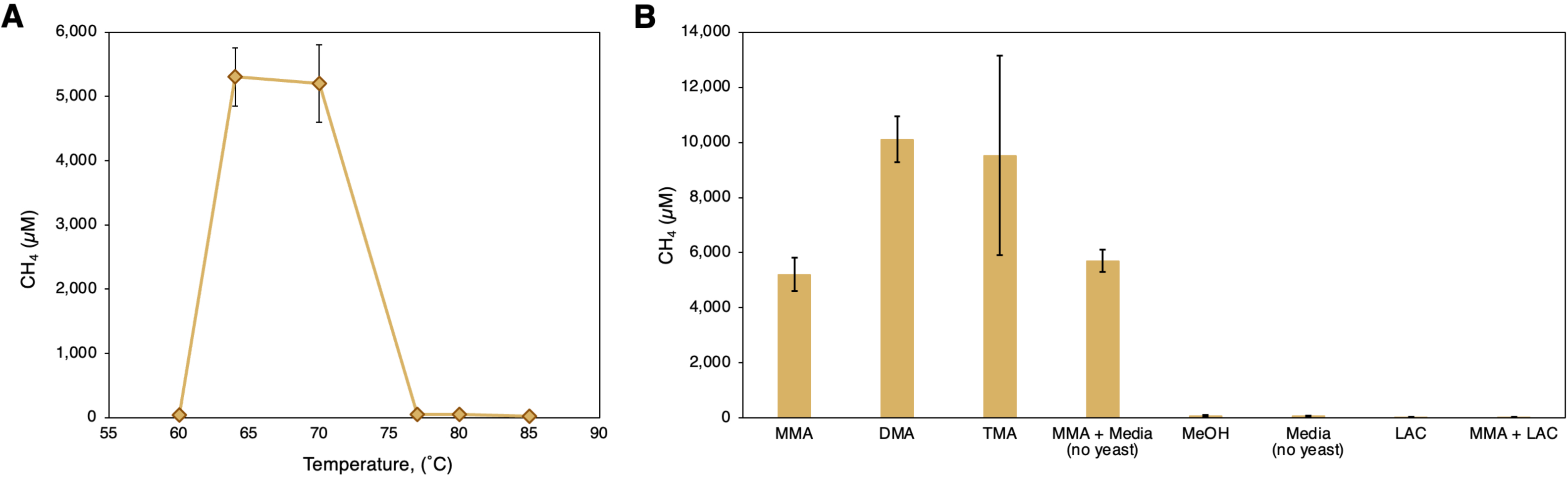
Temperature and substrate range of culture LCB24. *(A)* Methane production from MMA was observed between 64-70 °C. *(B)* Substrate range. Methane production was observed for MMA, DMA, TMA, and in media prepared without yeast extract. LAC, lactate; MeOH, methanol. Both experiments performed in triplicate. All measurements can be found in Dataset S4.

**Fig. 5.**
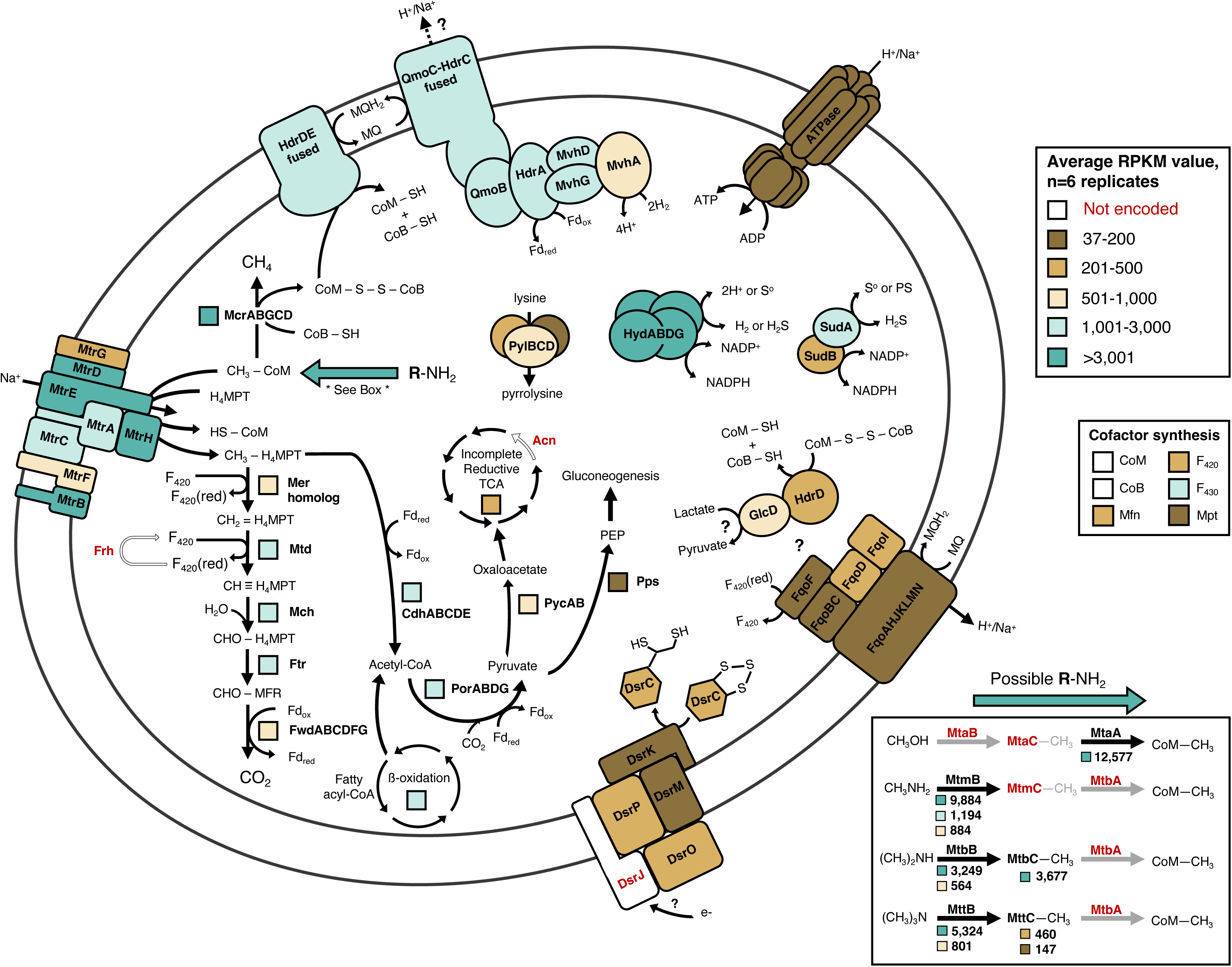
Transcriptional activity in *Ca.* M. hypatiae grown under methanogenic conditions (N_2_ headspace, 10 mM MMA, and 0.01% yeast extract). Transcriptionally active proteins are shown in bold black font. Proteins not encoded in the MAG are colored in white and denoted in bold red font. Average reads per kilobase of transcript per million mapped reads (RPKM) values of six biological replicates are depicted. RPKM values are represented by boxes or colored subunits close to each protein and are colored according to their expression level with the RPKM value of the lowest expressed gene depicted, 37 RPKM. For enzymes comprising multiple subunits, the beta-oxidation pathway, and the TCA cycle, an average RPKM value representing the transcribed enzymes is used. *Ca.* M. hypatiae is transcriptionally active under methanogenic conditions and encodes the ability to convert methyl-groups from mono-, di-, and trimethylamine to methane. This ability is enabled by several copies of substrate-specific methyltransferases and corrinoid proteins highlighted in the box to the bottom right. A complete list of genes described in this figure, their transcription levels, and their abbreviations is provided in Dataset S5.

*Ca.* M. hypatiae expressed the methyl-branch of the Wood-Ljungdahl pathway (WLP) and the acetyl-CoA decarbonylase/synthase complex (Cdh, *cdhABCDE*), which is consistent with genes observed and shown to be expressed in sulfate-reducing Archaeoglobi genomes (55). This includes two paralogous copies of 5,10-methylenetetrahydromethanopterin reductase (*mer*) which might function as a traditional Mer, considering that these genes are also members of the large luciferase-like monooxygenase family (pfam00296)(35). The expression of genes in the WLP varied. Methylenetetrahydromethanopterin dehydrogenase (*mtd*), methenyltetrahydromethanopterin cyclohydrolase (*mch*), formylmethanofuran-tetrahydromethanopterin N-formyltransferase (*ftr*), formylmethanofuran dehydrogenase (*fwdABC*), and one copy of the *mer* homologs were expressed at comparatively high levels (456-2,763 RPKM), whereas FwdDEFG and the other *mer* copy were only minimally expressed (<180 RPKM) (Dataset S5). The high expression of the Cdh complex (*cdhACDE*; 3,063±362, *cdhB* 677 RPKM average across subunits) suggests that *Ca.* M. hypatiae is capable of autotrophically fixing CO_2_ to acetyl-CoA as has been shown for other *Archaeoglobus* species (59). Acetyl-CoA could also be derived from the degradation of fatty acids present in yeast extract through the process of beta-oxidation. Enzymes involved in this pathway were expressed at moderate to high levels during growth (Dataset S6). Pyruvate synthase (Por) was highly expressed providing a way for acetyl-CoA to be converted to pyruvate and subsequently be fed into major biosynthetic pathways. Specifically, *Ca.* M. hypatiae encodes pyruvate carboxylase (PycAB), an incomplete reductive tricarboxylic acid cycle (rTCA), phosphoenolpyruvate synthase (Pps), most enzymes needed for gluconeogenesis, and several enzymes associated with the pentose phosphate pathway in archaea, which were all expressed at varying levels (Dataset S5). Together, these pathways provide *Ca.* M. hypatiae the capacity to synthesize amino acids, carbohydrates, integral components of the cell wall, and vital sugars for nucleic acids.

Several complexes related to energy conservation and electron transport were moderately to highly expressed. *Ca.* M. hypatiae encodes a fused heterodisulfide reductase (*hdrDE*) that was highly expressed (1,106±120 RPKM) in addition to a fused *hdrD/mvhD* and four copies of *hdrD* that were all expressed at much lower levels (<500 RPKM). The differing levels of transcription suggest the membrane-bound HdrDE is responsible for the regeneration of coenzymes M and B through the reduction of heterodisulfide (CoM-S-S-CoB). Additionally, the absence of HdrB, which contains the active site for disulfide reduction, eliminates the possibility that disulfide reduction could occur via a HdrABC complex (60). As reported for *Ca*. M. nevadensis (35), a unique gene cluster was identified containing F_420_-non-reducing hydrogenase (MvhAGD), two HdrA copies and a QmoC fused to a HdrC. One HdrA copy (JOOIALLP_01710) was predicted by DiSCo analysis as a quinone-modifying oxidoreductase (QmoB), a protein related to the HdrA of methanogens (61, 62). This cluster was expressed at high levels (995-2,431 RPKM average), suggesting its importance for electron transfer in *Ca.* M. hypatiae. We hypothesize these subunits are associating together *in vivo* to bifurcate electrons from hydrogen (H_2_) to reduce both menaquinone (MQ) and ferredoxin (Fd_ox_), as proposed recently (35, 63). Lastly, *Ca.* M. hypatiae moderately expressed a membrane-bound F_420_H_2_:quinone oxidoreductase (Fqo) complex (88-280 RPKM across subunits) and a V-type ATP synthase (24-442 RPKM across subunits).

The electrons required for reducing the CoM-S-S-CoB heterodisulfide could originate from two possible routes. The first possibility would rely on sourcing electrons from hydrogen, which could be oxidized by the Mvh-Qmo-Hdr complex coupled to menaquinone reduction. H_2_ may be produced through the activity of a group 3b [NiFe]-sulfhydrogenase (HydABDG), which was the highest expressed hydrogenase complex with an average RPKM of 4,421 across subunits (64, 65). To evolve hydrogen via HydABDG, reducing power, via NADPH, could be supplied by sulfide dehydrogenase (SudAB; SudA, 1,088 RPKM; SudB, 495 RPKM). Alternatively, NADPH could instead be provided to biosynthesis pathways and therefore be decoupled from methanogenic metabolism. H_2_ could also potentially be sourced from fermentative bacteria in the enrichment culture, however, the low number of hydrogenases encoded by co-enriched organisms were only very lowly expressed at the time of sampling for metatranscriptomics (<51 RPKM). At this point, the source of H_2_ *Ca.* M. hypatiae uses remains uncertain, as no H_2_ was added to the headspace. Second, in a hydrogen-independent electron transport system, reduced F_420_ and ferredoxin could be produced through the dismutation of methylated substrate to CO_2_ through the WLP. Reduced F_420_ could be oxidized by the Fqo complex and contribute to a reduced menaquinone pool that could be used by the fused HdrDE complex to reduce CoM-S-S-CoB. Reduced ferredoxin could be oxidized at a soluble FqoF to reduce F_420_ or at an Fqo complex lacking FqoF to reduce menaquinone (66, 67). Based on the low expression levels of the Fqo complex (171±67 RPKM) and the absence of F_420_-reducing hydrogenase (*frh*) from the genome, it is not likely the WLP runs in the reductive direction as a source of reduced F_420_ would be required. Resolving the exact configuration of the electron transport system encoded by *Ca.* M. hypatiae will require biochemical confirmation in future investigations.

Importantly, genes necessary for dissimilatory sulfate reduction typically observed in sulfate-reducing members of the Archaeoglobi, including dissimilatory sulfite reductase (*dsrAB*), sulfate adenylyltransferase (*sat*), and adenylylsulfate reductase (*aprAB*), were neither identified in the genome of *Ca.* M. hypatiae nor in the unbinned fraction of the metagenome. They were also absent from the comparatively incomplete MAG of *Ca.* M. nevadensis GBS (35). However, *Ca.* M. hypatiae encodes subunits *dsrMK* and *dsrOP* of the Dsr complex in addition to *dsrC*. This complex is strictly conserved in sulfate-reducing organisms (68) where it mediates electron transfer from the periplasm to the cytoplasm reducing the disulfide bond found in DsrC cysteines (69). The expression of the Dsr complex and *dsrC* was low (450±63 RPKM) during growth on monomethylamine suggesting it is not vital to the metabolism of *Ca.* M. hypatiae. The presence of the Dsr complex, DsrC, and subunits QmoC and QmoB in the genome may be explained as evolutionary remnants from ancestral Archaeoglobi, growing initially as sulfate-reducing organisms but later transitioning to a methanogenic lifestyle (7, 8). This raises the question whether intermediate of this process, Archaeoglobi capable of both methanogenesis and sulfate-reduction (and possible anaerobic oxidation of methane), still exist today (25, 28).

Collectively, the metagenomic and transcriptomic data confirmed that *Ca.* M. hypatiae is not only the sole archaeon but the sole methanogen in our culture. The metabolic reconstruction and metatranscriptomic results are consistent with methylotrophic methanogenesis from methylamines. The absence of genes required for sulfate reduction eliminates the possibility for this metabolism in *Ca.* M. hypatiae. A unique gene cluster (Mvh-Qmo-Hdr) potentially involved in energy conservation was expressed, however future studies will be required to test how *Ca.* M. hypatiae internally cycles electrons for methanogenesis and if it sources H_2_, or other reductants, from the medium or co-enriched bacteria.

### Distribution of *Ca.* Methanoglobus Across Geothermal Features in YNP

16S rRNA and *mcrA* gene amplicon sequence data generated in a recent microbial diversity survey of 100 geothermal features in YNP (33) were used to analyze the distribution of Archaeoglobi related to *Ca.* M. hypatiae (SI Appendix, Fig. S5). 16S rRNA gene amplicons closely related to *Ca.* M. hypatiae (96.7-100% sequence identity) were found in seven DNA samples from six hot springs (pH 5.1-9.35, 31-78 °C) in addition to hot spring LCB024 (the source of this culture) at relative abundances ranging from 0.02-0.22%. In addition, *mcrA* gene ASVs affiliated with Archaeoglobi were PCR-amplified from 53 DNA samples, out of 201 total samples that had been screened by PCR. These 53 samples had been collected from microbial mats or sediments originating from 36 geothermal features distributed across various thermal regions within YNP by Lynes, Krukenberg *et al.* (33). Archaeoglobi-related *mcrA* genes were found in geothermal features with a pH range of 2.61 to 9.32 and a temperature range of 18.4 °C to 93.8 °C. Collectively, our results and the studies by Wang *et al*. and Buessecker *et al*., who reported that Mcr-encoding Archaeoglobi are present (35) and transcriptionally active in hot spring mesocosms (34), demonstrate the previously overlooked role that Archaeoglobi might play in the anaerobic carbon cycle of geothermal environments.

## Conclusion

In summary, the cultivation of *Ca.* Methanoglobus hypatiae LCB24 provides direct experimental evidence that members of the Archaeoglobi are methanogens. *Ca.* M. hypatiae can use MMA, DMA, and TMA as methanogenic substrates and grows optimally at 64-70 °C, as evidenced by metagenomics, metatranscriptomics, and isotope tracing experiments. Metagenomic sequencing and phylogenomic analysis confirmed the close relationship of *Ca.* M. hypatiae to other Mcr-encoding Archaeoglobi and the relatedness of its *mcrA* to MAGs of the TACK superphylum, some of which have recently been shown to also be methanogens (70, 71). Together, this supports the idea that the capacity for methanogenesis is deeply rooted in the archaea and possibly dates to the last common ancestor of archaea (1, 3, 7, 8, 72). The wide distribution of Archaeoglobi-affiliated *mcrA* gene sequences and *Ca.* M. hypatiae-related 16S rRNA gene sequences in geothermal features across YNP suggests that members of this lineage play a hitherto unaccounted-for role in anaerobic carbon cycling in these extreme ecosystems. Future studies of *Ca.* M. hypatiae and other methanogens will provide valuable insights into the evolution of methane metabolism and the significance of these archaea in biogeochemical cycles across geothermal and other environments.

## Acknowledgements

This study was funded through a NASA Exobiology program award (80NSSC19K1633) to R.H. We thank the US National Park Service for permitting work in YNP under permit number YELL-SCI-8010. We thank George Schaible (MSU) for help with SEM imaging, Dr. Viola Krukenberg (MSU) for initial FISH methodology development, Sylvia Nupp, Dr. Andrew Montgomery, and Paige Schlegel (all MSU) for assistance with field sampling, Dr. Christopher Lemon (MSU) for allowing use of his cooling centrifuge, and Dr. Marike Palmer (UN Las Vegas) for discussing naming of this archaeon.

## Author Contributions

M.M.L. and R.H. developed the research project. M.M.L., Z.J.J., A.J.K, and R.H. designed experiments. M.M.L. and A.J.K. conducted field sampling. M.M.L. performed cultivation, extracted DNA for amplicon and metagenomic sequencing, extracted RNA for transcriptomic sequencing, and conducted physiology and stable isotope experiments. A.J.K. developed GC/GCMS protocols and processed GCMS samples. Z.J.J. processed and annotated metagenomic and transcriptomic data, assembled MAGs, mapped transcripts, assigned taxonomy, constructed 16S rRNA gene phylogeny, and performed phylogenetic analysis of MAGs. M.M.L. conducted phylogenetic analysis of amplicon data, refined gene annotations, reconstructed, and interpreted the metabolic potential of *Ca.* M. hypatiae with insight from Z.J.J and A.J.K. R.H. was responsible for funding and supervision of the project. M.M.L. and R.H. wrote the manuscript, which was then edited by all authors.

## Supporting Information

### SI Results and Discussion

#### Protologue

##### *Methanoglobus hypatiae* sp. nov

Me.tha.no.glo.bus. Gr. pref. *methano-*, pertaining to methane; L. masc. n. -*globus*, sphere; Gr.L. masc. n. *Methanoglobus*, methane producing organism spherical in shape. This genus was named by Buessecker *et al.* (1). Hy.pa.ti.ae. Gr. fem. hypatiae, to honor Hypatia of Alexandria, a respected and renowned philosopher of ancient Alexandria, Egypt, who made significant contributions to the understanding of mathematics and astronomy. A symbol of intellectual courage and scholarly achievement. This archaeon was cultured from an unnamed hot spring in the Lower Culex Basin of Yellowstone National Park identified as feature LCB024 (2). This archaeon is an obligately anaerobic thermophile that performs methylotrophic methanogenesis using methylamines and grows as regular to irregular coccoid cells approximately 0.5 to 1 µm in width. The type genome of this archaeon is deposited at NCBI under BioProject PRJNA1014417, accession number will be added upon publication.

### SI Materials and Methods

#### Amplicon Sequencing and Analysis

DNA was extracted from environmental slurry samples and enrichment cultures sampled on the day of transfer using the FastDNA Spin Kit for Soil (MP Biomedicals, Irvine, CA) following the manufacturer’s guidelines. Archaeal and bacterial 16S rRNA genes were amplified with the updated Earth Microbiome Project primer set 515F and 806R (3). Amplicon libraries were prepared as previously described (2) and sequenced by the Molecular Research Core Facility at Idaho State University (Pocatello, ID) using an Illumina MiSeq platform with 2 x 250 bp paired end read chemistry. Gene reads were processed using QIIME 2 version 2022.8 (4). Primer sequences were removed from demultiplexed reads using cutadapt (5) with error rate 0.12 and reads truncated (130 bp forward, 150 bp reverse), filtered, denoised and merged in DADA2 with default settings (6). Processed 16S rRNA gene amplicon sequence variants (ASVs) were taxonomically classified with the sklearn method and the SILVA 138 database (7). The R package *decontam* (version 1.18.0) (8) was used to remove contaminants using the “Prevalence” model with a threshold of 0.5.

### Annotation and Reconstruction of Metabolic Potential

Genes associated with methanogenesis pathways, dissimilatory sulfur metabolism pathways, coenzyme and cofactor biosynthesis, energy conservation, and beta-oxidation, were inventoried. Annotations assigned by Prokka were refined through manual evaluation using KofamKOALA, NCBI BLASTP, NCBI’s Conserved Domain Database, InterPro, the hydrogenase classifier HydDB, and DiSCo (9–14).

### Phylogenetic and Phylogenomic Analyses

Average nucleotide identities (ANI) of 16S rRNA genes were calculated with blastn, with ANI and average amino acid identities (AAI) calculated by pyani v02.2.12 (ANIb) and CompareM v0.0.23 (-- fragLen 2000) (https://github.com/dparks1134/CompareM), respectively for selected Archaeoglobales genomes and MAGs (Table 1). Phylogenetic analysis of 16S rRNA genes was performed with fasttree (15) using masked alignments generated by ssu-align.

Archaeoglobales MAGs and reference genomes were screened for 54 phylogenetically informative single copy proteins (16, 17) of which a subset of 33 proteins were identified in them all (SI Appendix, Table S4). In order to maximize the number of proteins compared across references, MAGs LMO1 and LMO3 were excluded from this analysis, as they lacked 2 and 7 proteins out of the total 33, respectively. These were then aligned with muscle (18), concatenated, and phylogenomically analyzed with maximum likelihood analysis with fasttree (WAG model). McrA alignments were performed with MAFFT-LINSi v7.522 (19), trimmed with trimAL v1.4.rev22 (20) using a 0.5 gap threshold, and maximum likelihood trees were built with IQTree2 v2.0.6 (21) using LG+C60+F+G model and 1,000 ultrafast bootstraps.

### Temperature and Substrate Optimum Experiments

Methane production and growth of Archaeoglobus was evaluated at different temperatures and in the presence of methylated substrates (i.e., methanol and mono-, di-, and trimethylamine), lactate, and media prepared without yeast extract. The sixth transfer of the enrichment was used to inoculate triplicate 30 mL serum bottles containing 15 mL of medium with 8% v/v inoculum, streptomycin (50 mg/L), vancomycin (50 mg/L), and 10 mM of each substrate tested. Cultures were evaluated at 60 °C, 64 °C, 70 °C, 77 °C, 80 °C, and 85 °C with 10 mM MMA. Separately, we tested whether the culture would grow on the following substrate (combinations): 10 mM dimethylamine (DMA); 10 mM trimethylamine (TMA); 10 mM methanol (MeOH); 10 mM lactate (LAC); 10 mM MMA and 10 mM LAC); 10 mM MMA with media without yeast extract; and a control in media without yeast or any methanogenic substrate. The 70 °C cultures amended with 10 mM MMA served as the control. All incubations were performed in biological triplicate.

### Methane Measurements

During cultivation, 250 µL subsamples of the headspace were taken using a gas tight syringe (Hamilton) and injected into a 10 mL autosampler vial that had been sealed with grey chlorobutyl septa. Samples were taken from the autosampler vials and injected into a Shimadzu 2020-GC gas chromatograph equipped with a GS-CarbonPLOT column (30 m x 0.32 mm; 1.5 μm film thickness; Agilent) and a Rt-Q-BOND column (30 m x 0.32 mm; 1.5 μm film thickness; Restek) using helium as a carrier gas. All injections were performed by a Shimadzu AOC-6000 autosampler robot. The injector, column, and flame ionization detector (FID) were maintained at 200 °C, 50 °C, and 240 °C, respectively. Methane concentrations were calculated based on injection of a standard curve.

### Fluorescence *in situ* hybridization and cell counts

Aliquots of enrichment cultures incubated with ^13^C-MMA during the SIT experiment were treated with 2% paraformaldehyde (PFA) and fixed for 1 hr at room temperature. Following fixation, cells were washed twice with 1x PBS, followed by centrifugation at 16,000 × g to remove the supernatant, resuspended in 1x PBS, and stored at 4 °C. For direct cell counts, aliquots of fixed cell suspensions were filtered onto polycarbonate filters (0.2 μm pore size, 25 mm diameter, GTTP Millipore, Germany) and air dried before filter pieces were cut and embedded in 0.2% low melting agarose. We attempted to use the Archaeoglobales-specific probe Arglo32 (22), however fluorescent signal was insufficient. Given *Ca.* M. hypatiae was the sole archaeon in the enrichment culture, the relative abundance of *Ca.* M. hypatiae cells was determined via catalyzed reporter deposition fluorescence *in situ* hybridization (CARD-FISH) using the general archaea-targeted 16S rRNA oligonucleotide probe Arch915 (23). Total cell counts were based on DNA-stained cells using DAPI (4,6-diamidino-2-phenylindole). CARD-FISH was performed as previously described (24). Cell wall permeabilization was achieved with a brief treatment of 0.1 M HCl (1 min, RT) followed by treatment with 0.01 M HCl (15 min, RT). Endogenous peroxidases were inactivated with 0.15% H_2_O_2_ in methanol (30 min, RT). A formamide concentration of 35% was used for all hybridization reactions (2.5 hrs, 46°C). CARD was performed using Alexa Fluor 594 labeled tyramides for 30 min at 46°C. Following signal amplification, an additional washing step in 1x PBS was included to reduce background fluorescence (15 min, RT, dark). Samples were stained with DAPI, embedded in Citifluor-Vectashield, and enumerated using an epifluorescence microscope (Leica DM4B).

### Scanning electron microscopy (SEM)

An aliquot of the enrichment culture at transfer 7 (T7) was treated with 2% paraformaldehyde (PFA) and fixed for 1 hr at room temperature. Following fixation, cells were washed twice through centrifugation at 16,000 × g to remove the supernatant, resuspended in 1x phosphate buffered saline (PBS), and stored at 4 °C. Samples for imaging were prepared according to Schaible et al., 2022 (25). Briefly, a square coupon of mirror-finished 304 stainless steel (25 mm diameter, 0.6 mm thickness) was purchased from Stainless Supply (Monroe, NC). The coupon was cleaned by washing with a 1% solution of Tergazyme (Alconox, New York, NY) and rinsed with Milli-Q water. The coupon was dried under compressed air and stored at room temperature. 5 µL of fixed sample was spotted on the coupon and air-dried at 46 °C for 3 min. The coupon was then dried for 1 m each step in a successive ethanol series starting with 10% ethanol and increasing by increments of 10% with the last step 90% ethanol. SEM images were captured using a Zeiss (Jena, Germany) SUPRA 55VP field emission scanning electron microscope (FE-SEM). The microscope was operated at 1 keV under a vacuum of 0.2–0.3 mPa, with a working distance of 5.4-6.2 mm at the Imagining and Chemical Analysis Laboratory (ICAL) of Montana State University (Bozeman, MT). No conductivity coating was applied before SEM analysis as the microscope was operated at 1 keV.

### SI Tables

**Table S1.** Extended community composition history of methanogenic enrichment cultures via estimated relative abundance (%) from 16S rRNA gene amplicon sequencing.

|  | LCB024<br>sediment | Slurry<br>11/2021 | Slurry<br>02/2022 | Initial<br>Enrichment | T1 | T2 | T3 | T4 | T4 -<br>MG | T5 | SIT-MG |
| --- | --- | --- | --- | --- | --- | --- | --- | --- | --- | --- | --- |
| Archaeoglobaceae<br>ASV_78ad2 | 0.46 | 0.32 | 0.02 | 6.42 | 74.8 | 68.9 | 48.5 | 46.0 | 84.7 | 62.2 | 92.8 |
| Pseudothermotoga<br>ASV_17231 | 2.14 | 1.06 | 1.08 | 18.93 | 6.8 | 17.1 | 28.5 | 37.4 | 14.5 | 15.7 | 3.2 |
| Other<br>Archaeoglobaceae<br>uncultured | 0.00 | 0.00 | 0.00 | 1.54 | 0.0 | 0.01 | 0.0 | 0.0 | 0.0 | 0.0 | 0.0 |
| Confirmed<br>methanogenic<br>archaea | 0.25 | 0.38 | 0.45 | 0.00 | 0.0 | 0.0 | 0.0 | 0.0 | 0.0 | 0.0 | 0.0 |
| Other Archaea | 13.51 | 11.87 | 10.49 | 2.67 | 0.0 | 0.0 | 0.0 | 0.04 | 0.0 | 0.0 | 0.0 |
| Other Bacteria | 83.63 | 86.37 | 87.95 | 70.44 | 18.4 | 14.1 | 23.1 | 16.5 | 0.8 | 22.1 | 4.0 |

**Table S2.** Methane production of the enrichment culture over time. T, transfer.

| Culture ID | Days continuous | Days | CH<sub>4</sub> (μM) | CH<sub>4</sub> (mM) | CH<sub>4</sub> (ppm) | CH<sub>4</sub> (%) |
| --- | --- | --- | --- | --- | --- | --- |
| **1A – initial enrichment** | 6 | 6 | 4.91 | 0.00 | 118.67 | 0.01 |
|  | 13 | 13 | 16.73 | 0.02 | 404.73 | 0.04 |
|  | 20 | 20 | 12.98 | 0.01 | 314.13 | 0.03 |
|  | 27 | 27 | 12.86 | 0.01 | 311.20 | 0.03 |
|  | 36 | 36 | 32.92 | 0.03 | 796.48 | 0.08 |
|  | 48 | 48 | 146.28 | 0.15 | 3538.80 | 0.35 |
|  | 53 | 53 | 279.15 | 0.28 | 6753.17 | 0.68 |
|  | 63 | 63 | 419.11 | 0.42 | 10139.33 | 1.01 |
|  | 70 | 70 | 447.12 | 0.45 | 10816.36 | 1.08 |
| **T1** | 82 | 12 | 14.40 | 0.01 | 348.40 | 0.03 |
|  | 91 | 21 | 50.85 | 0.05 | 1230.06 | 0.12 |
|  | 98 | 27 | 198.41 | 0.20 | 4800.09 | 0.48 |
|  | 106 | 35 | 390.47 | 0.39 | 9445.97 | 0.94 |
|  | 113 | 42 | 565.32 | 0.57 | 13676.04 | 1.37 |
|  | 117 | 46 | 688.44 | 0.69 | 16654.06 | 1.67 |
|  | 126 | 55 | 1442.03 | 1.44 | 34885.74 | 3.49 |
|  | 133 | 62 | 1844.13 | 1.84 | 44611.75 | 4.46 |
| **T2** | 146 | 7 | 15.22 | 0.02 | 368.32 | 0.04 |
|  | 158 | 18 | 16.11 | 0.02 | 389.83 | 0.04 |
|  | 175 | 34 | 49.40 | 0.05 | 1195.06 | 0.12 |
|  | 197 | 57 | 680.54 | 0.68 | 16463.41 | 1.65 |
|  | 221 | 81 | 2200.24 | 2.20 | 53227.90 | 5.32 |
|  | 229 | 89 | 2654.34 | 2.65 | 64213.66 | 6.42 |
|  | 238 | 98 | 2230.72 | 2.23 | 53964.66 | 5.40 |
| **T3** | 243 | 5 | 18.17 | 0.02 | 439.53 | 0.04 |
|  | 259 | 21 | 10.70 | 0.01 | 258.82 | 0.03 |
|  | 266 | 28 | 536.13 | 0.54 | 12969.71 | 1.30 |
|  | 274 | 36 | 2413.23 | 2.41 | 58379.47 | 5.84 |
|  | 280 | 42 | 2283.95 | 2.28 | 55253.34 | 5.53 |
|  | 284 | 46 | 2193.96 | 2.19 | 53077.04 | 5.31 |
| **T4** | 294 | 10 | 17.36 | 0.02 | 420.07 | 0.04 |
|  | 301 | 17 | 18.12 | 0.02 | 438.37 | 0.04 |
|  | 309 | 25 | 12.51 | 0.01 | 302.54 | 0.03 |
|  | 319 | 35 | 1944.01 | 1.94 | 47028.08 | 4.70 |
|  | 327 | 43 | 3054.00 | 3.05 | 73881.94 | 7.39 |
|  | 335 | 51 | 2619.45 | 2.62 | 63370.12 | 6.34 |
| **T5** | 347 | 10 | 13.41 | 0.01 | 324.42 | 0.03 |
|  | 355 | 18 | 13.14 | 0.01 | 317.78 | 0.03 |
|  | 365 | 28 | 137.59 | 0.14 | 3328.60 | 0.33 |
|  | 371 | 34 | 1023.35 | 1.02 | 24756.27 | 2.48 |
|  | 375 | 38 | 2218.75 | 2.22 | 53674.81 | 5.37 |
|  | 384 | 46 | 2459.29 | 2.46 | 59493.85 | 5.95 |
| **T6** | 386 | 2 | 16.29 | 0.02 | 394.19 | 0.04 |
|  | 393 | 9 | 21.33 | 0.02 | 516.14 | 0.05 |
|  | 400 | 16 | 18.68 | 0.02 | 451.92 | 0.05 |
|  | 407 | 23 | 245.86 | 0.25 | 5947.76 | 0.59 |
|  | 412 | 28 | 2555.27 | 2.56 | 61815.61 | 6.18 |
|  | 414 | 30 | 3432.93 | 3.43 | 83051.24 | 8.31 |
|  | 416 | 32 | 3621.90 | 3.62 | 87623.79 | 8.76 |
|  | 418 | 34 | 3942.97 | 3.94 | 95385.88 | 9.54 |
| **T7** | 428 | 9 | 34.33 | 0.03 | 830.38 | 0.08 |
|  | 435 | 16 | 296.91 | 0.30 | 7182.55 | 0.72 |
|  | 440 | 21 | 2931.05 | 2.93 | 70907.06 | 7.09 |
|  | 444 | 25 | 3164.45 | 3.16 | 76553.52 | 7.66 |
|  | 446 | 27 | 3338.04 | 3.34 | 80751.55 | 8.08 |

**Table S3.** Extended Archaeoglobales metagenome assembled genome and isolate genome statistics. A combined assembly of metagenomes from T4-MG and SIT-MG (Fig. 1A) for the *Ca.* M. hypatiae LCB24 MAG was used as it yielded an improved assembly. GTDB classified the YNP, GBS, and China MAGs as: d_Archaea;p_Halobacteriota;c_Archaeoglobi;o_Archaeoglobales; f_Archaeoglobaceae;g_WYZ-LMO2;s_WYZ-LMO2. Len., length; Compl., completeness; Redun, redundancy; Strain Hetero., strain heterogeneity; pOGT, predicted optimal growth temperature. * stop codon interrupts *mcrA* sequence; ^a^ Both sequences 5 start; not identical; ^b^ Consists of 1 chromosome and 1 plasmid.

|  | Seqs | Len (Mb) | GC (%) | Compl. (%) | Redun. (%) | 16S | tRNA | CDS | *mcrA* | CRISPRs | pOGT (°C) | Citation |
| --- | --- | --- | --- | --- | --- | --- | --- | --- | --- | --- | --- | --- |
| ***Ca. M. hyptiae* LCB24** | 19 | 1.624 | 46.12 | 100 | 1.31 | 1 | 44 | 1,760 | 1 | 2 | 74.44 | This study |
| Archaeoglobales LCB024-003 | 179 | 1.157 | 45.72 | 88.48 | 1.31 | 0 | 31 | 1,226 | 1 | 1 | 73.90 | Lynes *et al.* 2023 |
| Archaeoglobales WYZ-LMO2 | 220 | 1.514 | 45.92 | 97.60 | 0 | 0 | 40 | 1,614 | 1 | 2 | 72.40 | Wang *et al.* 2019 |
| *Ca. M. hydrogenotrophicum* bin74 | 127 | 1.547 | 45.63 | 92.81 | 1.31 | 1 | 34 | 1,679 | 0* | 9 | 75.32 | Liu *et al.* 2020 |
| Archaeoglobales SJ3.Bin34 | 73 | 1.468 | 46.00 | 98.69 | 0.65 | 0 | 39 | 1,581 | 1 | 2 | 73.13 | Colman *et al.* 2019 |
| *Ca. Methanoglobus nevadensis* GBS | 71 | 1.602 | 47.16 | 98.04 | 3.66 | 1 | 47 | 1,785 | 1 | 2 | 74.75 | Peacock *et al.* 2013 |
| Archaeoglobales GMQP bin32 | 20 | 1.727 | 41.26 | 100 | 0.98 | 1 | 39 | 1,931 | 1 | 2 | 71.88 | Hua *et al.* 2019 |
| Archaeoglobales GMQP_D bin 18 | 22 | 1.565 | 42.01 | 97.39 | 0.98 | 1 | 45 | 1,747 | 1 | 3 | 69.41 | Wang *et al.* 2023 |
| Archaeoglobales JZ-3 D bin 138 | 32 | 1.550 | 42.08 | 99.35 | 1.31 | 1 | 43 | 1,701 | 1 | 2 | 69.11 | Wang *et al.* 2023 |
| Archaeoglobales WYZ-LMO1 | 140 | 1.557 | 43.85 | 88.89 | 1.31 | 1 | 41 | 1,701 | 1 | 1 | 72.72 | Wang *et al.* 2019 |
| Archaeoglobales WYZ-LMO3 | 135 | 1.568 | 43.93 | 88.03 | 1.96 | 2<sup>a</sup> | 35 | 1,670 | 1 | 2 | 73.09 | Wang *et al.* 2019 |
| *Ca. M. hydrogenotrophicum* bin11 | 252 | 1.220 | 47.59 | 91.83 | 6.17 | 1 | 27 | 1,387 | 1 | 0 | 72.64 | Liu *et al.* 2020 |
| *Ca. M. hydrogenotrophicum* bin16 | 46 | 1.668 | 45.49 | 96.51 | 1.31 | 0 | 41 | 1,864 | 1 | 2 | 72.69 | Liu *et al.* 2020 |
| Archaeoglobus JZ bin24 | 35 | 1.492 | 44.66 | 99.35 | 1.31 | 0 | 44 | 1,642 | 0 | 2 | 68.52 | unpublished |
| Archaeoglobales LCB024-002 | 66 | 1.334 | 42.28 | 97.39 | 0.03 | 0 | 41 | 1,459 | 0 | 0 | 73.46 | Lynes *et al.* 2023 |
| Archaeoglobales LCB003-04 | 128 | 1.335 | 42.27 | 96.73 | 1.96 | 0 | 38 | 1,447 | 0 | 0 | 73.71 | Lynes *et al.* 2023 |
| Archaeoglobales SJ3.bin61 | 45 | 1.270 | 41.68 | 89.54 | 0 | 1 | 40 | 1,392 | 0 | 0 | 73.34 | Colman *et al.* 2019 |
| Archaeoglobales JZ_75 SW bin109 | 16 | 1.562 | 39.48 | 99.35 | 0.65 | 1 | 45 | 1,762 | 1 | 5 | 74.34 | Wang *et al.* 2023 |
| *Archaeoglobus fulgidus* DSM 4304 | 1 | 2.178 | 48.58 | 100 | 0 | 1 | 46 | 2,440 | 0 | 3 | 79.88 | Klenk *et al.* 1997 |
| *Archaeoglobus neptunius* SE56 | 32 | 2.116 | 46.05 | 100 | 0 | 1 | 47 | 2,336 | 0 | 2 | 78.26 | Slobodkina *et al.* 2021 |
| *Archaeoglobus profundus* DSM 5631 | 2<sup>b</sup> | 1.563 | 46.05 | 100 | 0 | 1 | 48 | 1,784 | 0 | 0 | 85.86 | von Jan *et al.* 2010 |
| *Ferroglobus placidus* DSM 10642 | 1 | 2.196 | 44.14 | 100 | 0 | 1 | 49 | 2,467 | 0 | 1 | 83.83 | Anderson *et al.* 2011 |
| *Geoglobus acetivorans* SBH6 | 1 | 1.861 | 46.84 | 99.84 | 0 | 1 | 48 | 2,168 | 0 | 6 | 78.22 | Mardanov *et al.* 2015 |
| *Geoglobus ahangari* 234 | 1 | 1.770 | 53.11 | 100 | 0 | 1 | 46 | 1,985 | 0 | 7 | 84.11 | Manzella *et al.* 2015 |
| *Archaeoglobus sulfaticallidus* PM70-1 | 1 | 2.077 | 43.24 | 100 | 0 | 1 | 51 | 2,237 | 0 | 1 | 76.12 | Stokke *et al.* 2013 |
| *Archaeoglobus veneficus* SNP6 | 1 | 1.902 | 47.05 | 99.35 | 0 | 1 | 46 | 2,055 | 0 | 2 | 76.18 | Mukherjee *et al.* 2017 |
| *Ca. Polytropus marinifundus* rG16 | 21 | 2.129 | 40.22 | 99.84 | 1.96 | 1 | 44 | 2,286 | 2 | 0 | 66.39 | Boyd *et al.* 2019 |

**Table S4.** Conserved single copy proteins used in phylogenomic analysis of MAGs and isolates.

| arCOG | Gene | Product |
| --- | --- | --- |
| arCOG00405 | GRS1 | Glycyl-tRNA synthetase (class II) |
| arCOG00779 | RplO | Ribosomal protein L15 |
| arCOG00785 | RpmC | Ribosomal protein L29 |
| arCOG01001 | Map | Methionine aminopeptidase |
| arCOG01183 | Kae1p/TsaD | Subunit of KEOPS complex, contains a domain with ASKHA fold and RIO-type kinase (AP-endonuclease activity) |
| arCOG01228 | Ffh | Signal recognition particle GTPase |
| arCOG01722 | RpsM/rps13p | Ribosomal protein S13 |
| arCOG01758 | RpsJ/rps10p | Ribosomal protein S10 |
| arCOG04070 | RplC | Ribosomal protein L3 |
| arCOG04071 | RplD | Ribosomal protein L4 |
| arCOG04072 | RplW | Ribosomal protein L23 |
| arCOG04086 | RpmD | Ribosomal protein L30 |
| arCOG04087 | RpsE | Ribosomal protein S5 |
| arCOG04088 | RplR | Ribosomal protein L18 |
| arCOG04090 | RplF/rpl6p | Ribosomal protein L6P |
| arCOG04091 | RpsH/rps8p | Ribosomal protein S8 |
| arCOG04094 | RplX/rpl24p | Ribosomal protein L24 |
| arCOG04095 | RplN/rps14p | Ribosomal protein L14 |
| arCOG04096 | RpsQ/rps17p | Ribosomal protein S17 |
| arCOG04097 | RpsC/rps3p | Ribosomal protein S3 |
| arCOG04098 | RplV/rpl22p | Ribosomal protein L22 |
| arCOG04113 | RplP | Ribosomal protein L10AE/L16 |
| arCOG04121 | RnhB | Ribonuclease HII |
| arCOG04169 | SecY | Preprotein translocase subunit SecY |
| arCOG04185 | RpsO | Ribosomal protein S15P |
| arCOG04239 | RpsD/rps4p | Ribosomal protein S4 or related protein |
| arCOG04242 | RplM/rpl13p | Ribosomal protein L13 |
| arCOG04243 | RpsI/rps9p | Ribosomal protein S9 |
| arCOG04245 | RpsB/rps2p | Ribosomal protein S2 |
| arCOG04255 | RpsL/rps12p | Ribosomal protein S12 |
| arCOG04256 | RpoC/Rpo11 | DNA-directed RNA polymerase subunit A" |
| arCOG04257 | RpoC/Rpo3/rpoA1 | DNA-directed RNA polymerase subunit A' |
| arCOG04277 | Efp | Translation elongation factor P (EF-P)/translation initiation factor 5A (eIF-5A) |

**Table S5.** Calculated cell density of replicates in the SIT experiment. FID measurements of replicates used to determine cell density. Density is calculated based on cell counts of DAPI and CARD-FISH labeled samples. Stdev., standard deviation. Letters A-F identify each replicate.

| Summary | <sup>12</sup>CH<sub>4</sub> |  | Cell Density, cells mL<sup>-1</sup> |  |  |
| --- | --- | --- | --- | --- | --- |
| Replicate ID | μM | ppm | DAPI | FISH | % labeled with Arch915 |
| 5A Day 22 | 25 | 610 | $3.83 \times 10^7$ | $1.34 \times 10^6$ | 3.5 |
| 5B Day 22 | 57 | 1,374 | $1.88 \times 10^7$ | $5.07 \times 10^6$ | 12.7 |
| 5E Day 22 | 50 | 1,204 | $3.49 \times 10^7$ | 0 | 0 |
| 5F Day 22 | 132 | 3,196 | $4.59 \times 10^7$ | $2.21 \times 10^6$ | 4.8 |
| **Day 22 Average** | 66 | 1,596 | $3.45 \times 10^7$ | $2.16 \times 10^6$ | 5.3 |
| **Day 22 Stdev.** | 46 | 1,116 | $1.14 \times 10^7$ | $2.15 \times 10^6$ | 5.4 |
| 5A Day 32 | 741 | 17,917 | $4.26 \times 10^7$ | $2.85 \times 10^7$ | 66.9 |
| 5B Day 32 | 2,176 | 52,635 | $1.11 \times 10^8$ | $6.00 \times 10^7$ | 54.2 |
| 5E Day 32 | 1,779 | 43,035 | $3.40 \times 10^7$ | $1.74 \times 10^7$ | 51.2 |
| 5F Day 32 | 2,412 | 58,355 | $9.14 \times 10^7$ | $4.03 \times 10^7$ | 44.1 |
| **Day 32 Average** | 1,777 | 42,985 | $6.97 \times 10^7$ | $3.65 \times 10^7$ | 54.1 |
| **Day 32 Stdev.** | 739 | 17,868 | $3.73 \times 10^7$ | $1.82 \times 10^7$ | 9.6 |
| 5A Day 45 | 4,109 | 99,398 | $12.2 \times 10^7$ | $6.41 \times 10^7$ | 52.7 |

### SI Figures

**Fig. S1.**
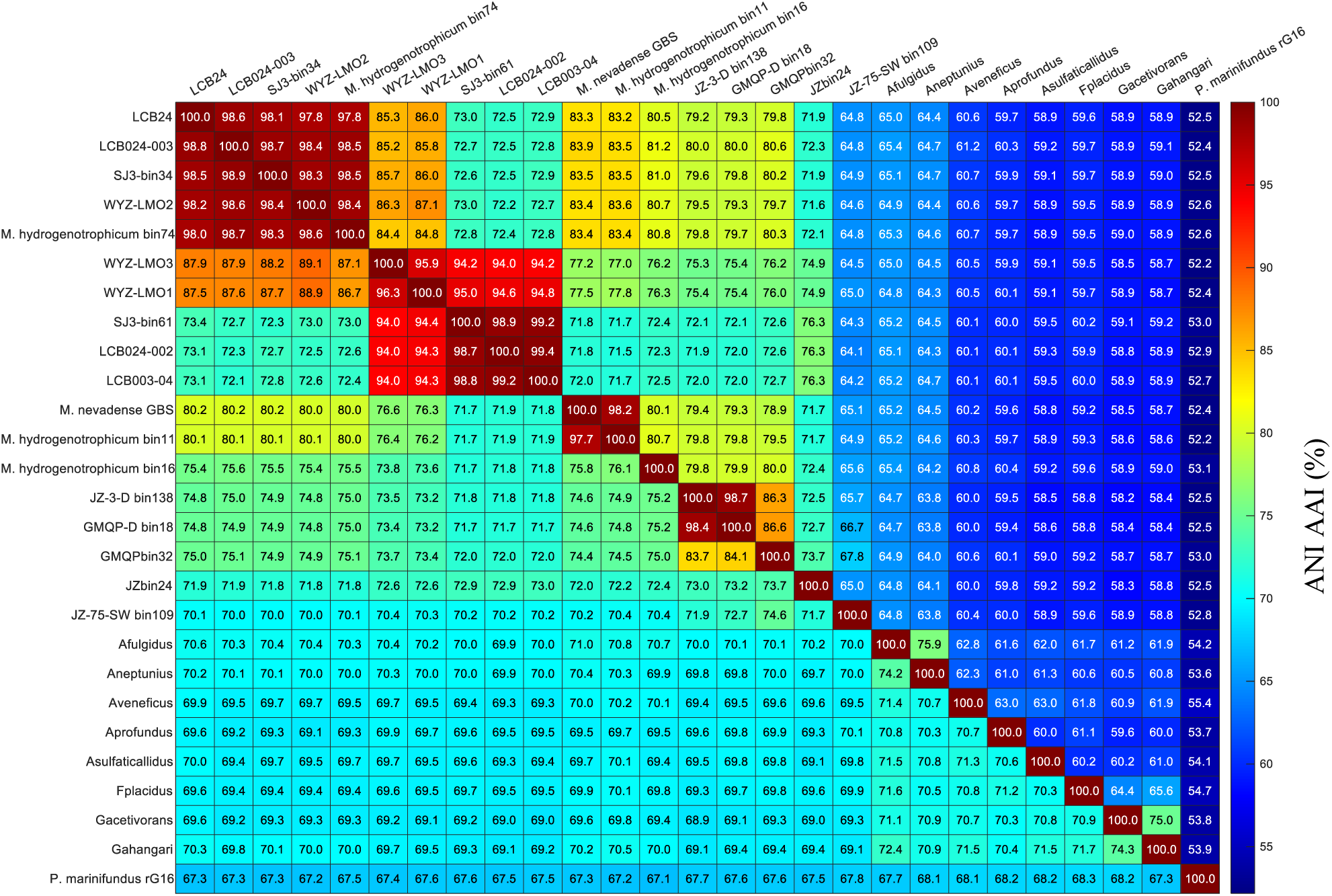
Detailed ANI (lower half of matrix) and AAI (upper half of matrix) analysis of related Archaeoglobales MAGs and reference genomes.

**Fig. S2.**
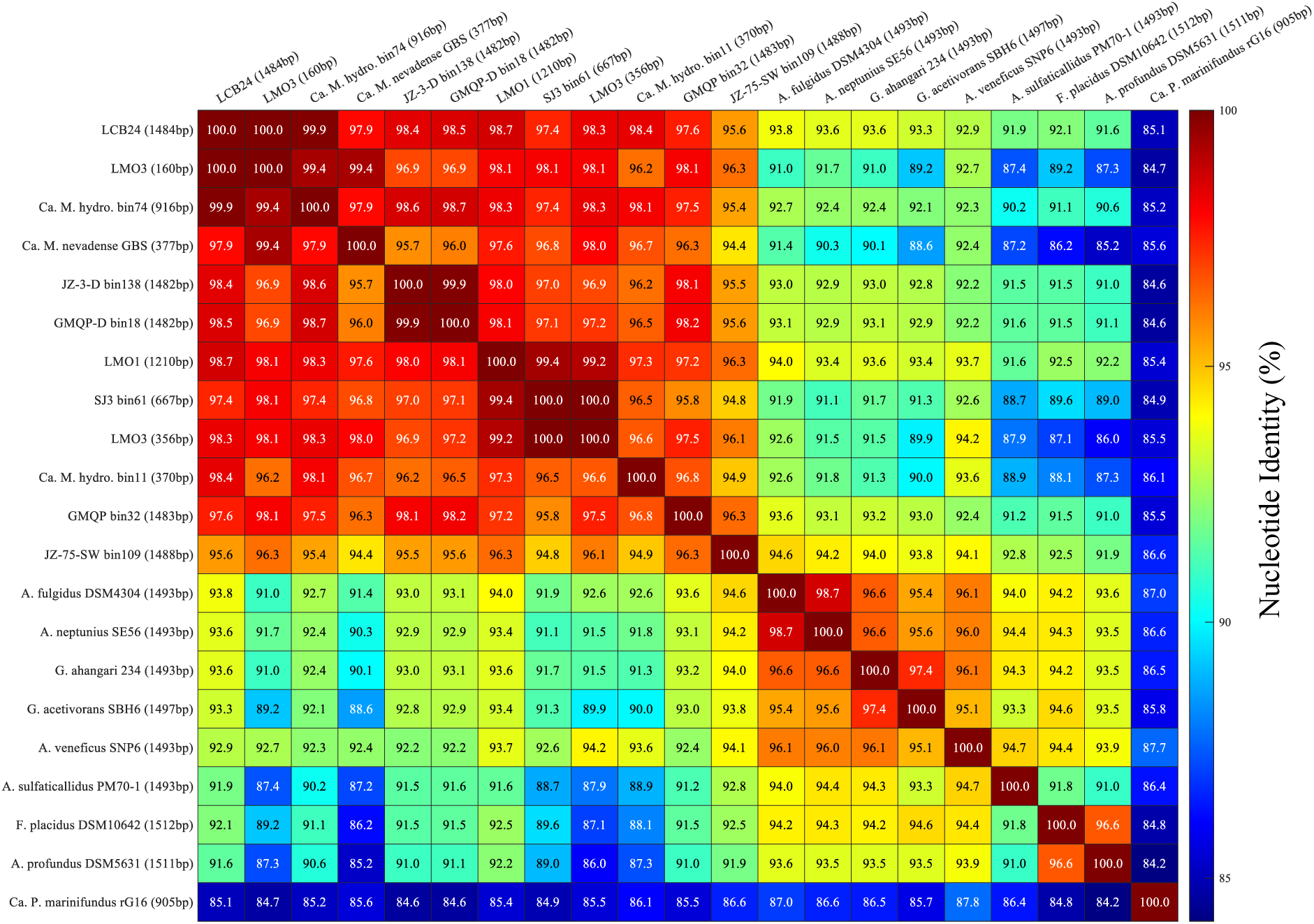
16S rRNA nucleotide identity analysis of closely related Archaeoglobales MAGs and reference genomes.

**Fig. S3.**
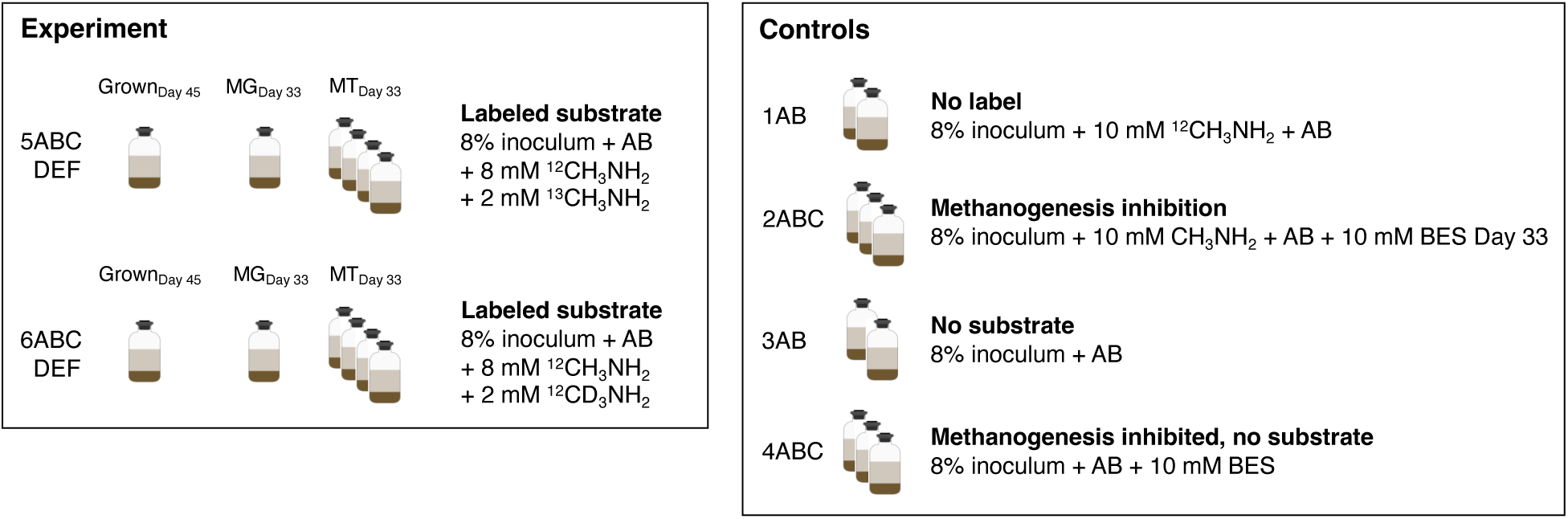
Experimental setup of the stable isotope tracing (SIT) experiment. Incubations were carried out in 30 mL culture volumes in 60 mL serum bottles with 8% v/v inoculum, 50 mg/L streptomycin, 50 mg/L vancomycin, 10 mM MMA, and N_2_ gas (99.999%) incubated in anoxic media (pH 7.8, 70 °C). Replicates sacrificed for analysis during mid-log phase are indicated. Of the eight samples harvested for metatranscriptomics, six were sequenced and used for analysis as two replicates did not yield sufficient RNA for sequencing. AB, antibiotics streptomycin and vancomycin, MG, metagenome sample; MT, metatranscriptome sample; BES, bromoethanesulfonate/methanogenesis inhibitor.

**Fig. S4.**
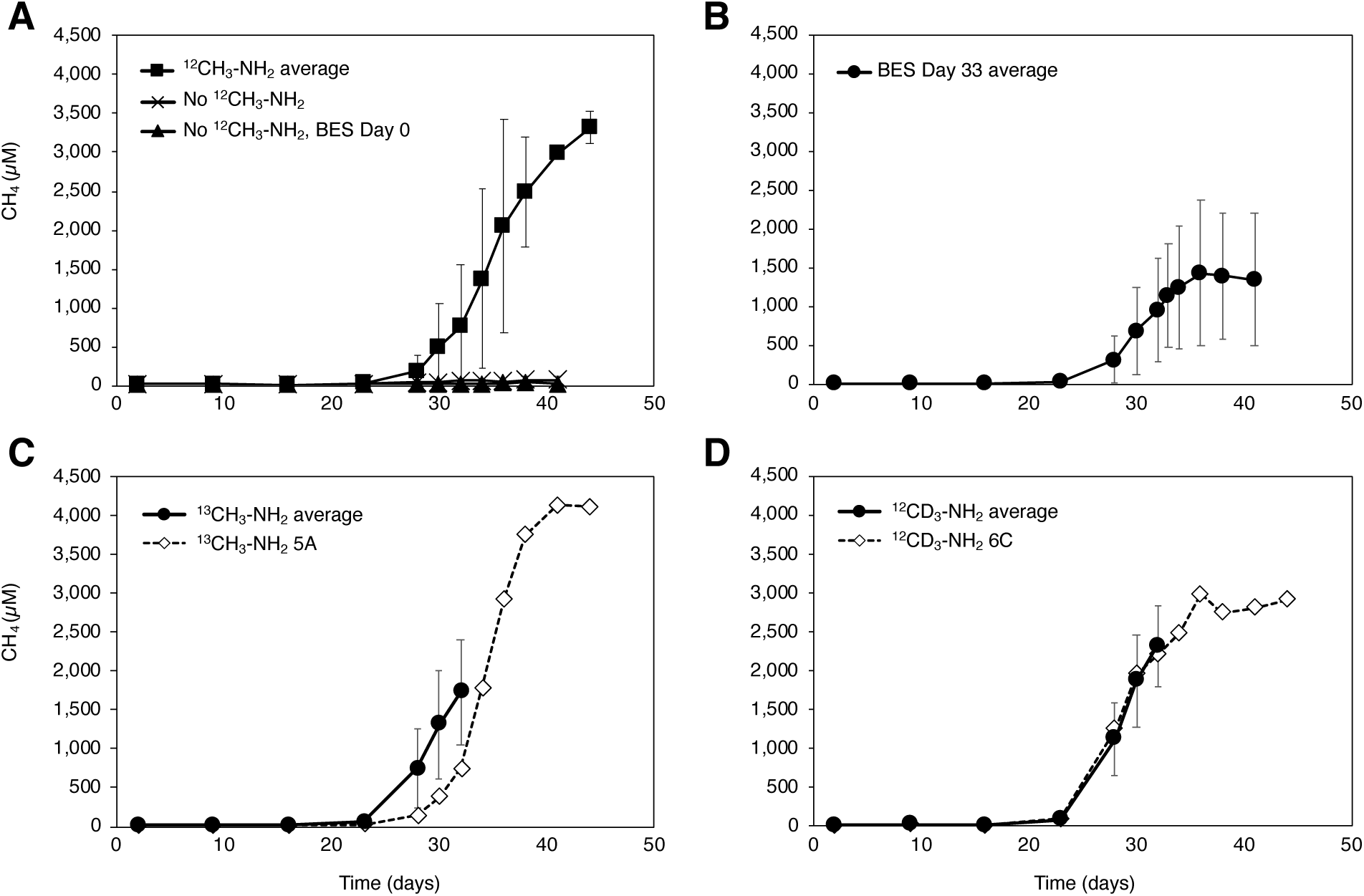
^12^CH_4_ measurements by GC-FID during the stable isotope tracing experiment. Measurements can be found in Dataset S3.

**Fig. S5.**
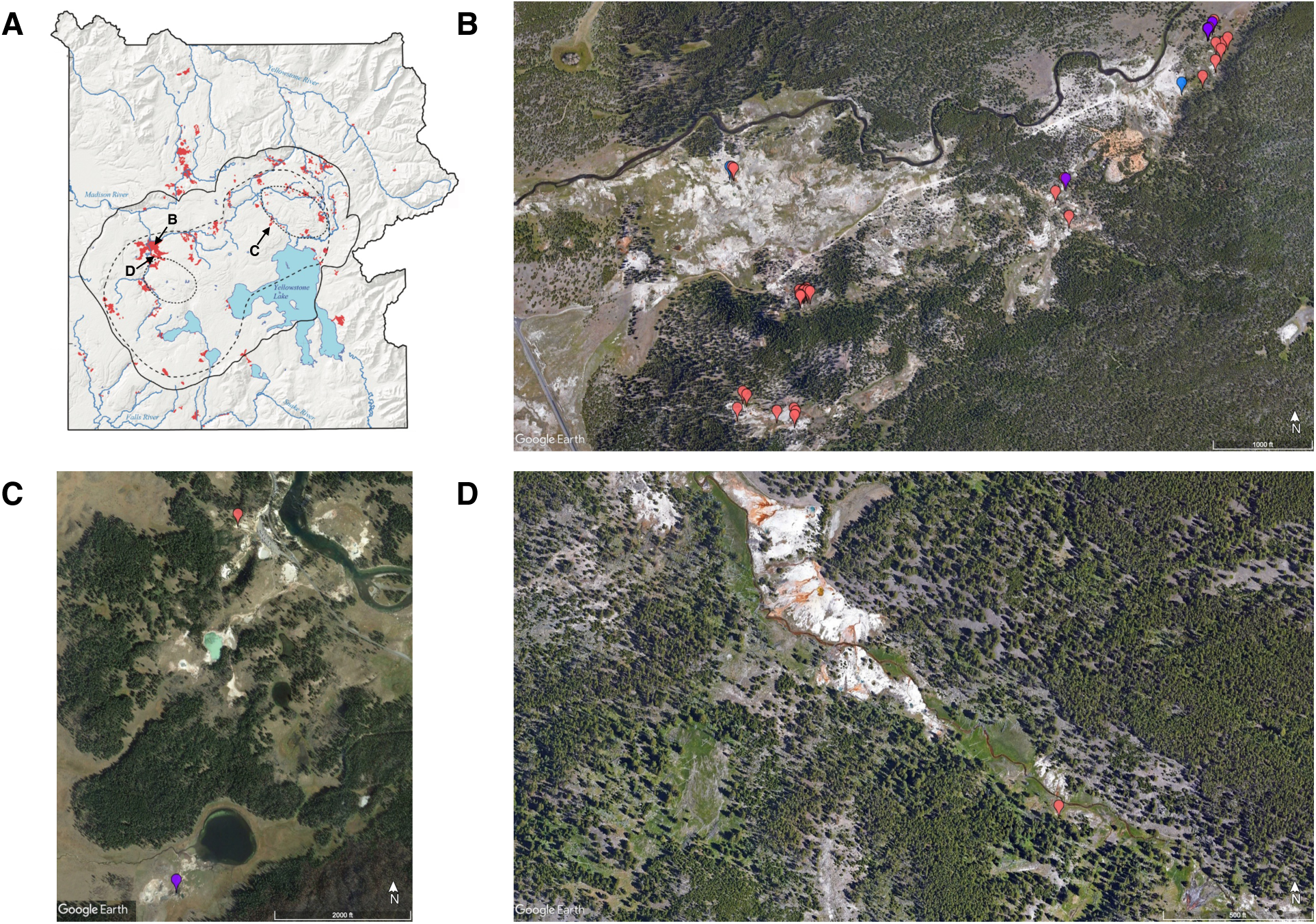
Geographical distribution of geothermal features in Yellowstone National Park in which Archaeoglobi-related *mcrA* genes (n = 36) and *Ca.* M. hypatiae-related 16S rRNA genes (n = 6) were detected. Features are located in the (*A*) Map of Yellowstone National Park Wyoming, USA modified from Vaughan *et al.* 2014 (26) *(B)* Lower Culex Basin (n = 36), (*C*) Mud Volcano Region (n = 2), and (*D*) the White Creek Area (n = 1). These features spanned a wide pH (2.61-9.35) and temperature (18.4-93.8 °C) range. Features in which *mcrA* were detected are marked in red and features with related 16S rRNA genes are shown in blue. Features in which both amplicons were detected are colored in purple. For details on these sites, their *mcrA* data, water geochemistry, and exact location, see Lynes & Krukenberg et al., 2023. Image source: Google Earth.

### Description of Available Supplementary Datasets

**SI Dataset S1.** Extended metagenome assembled genome (SIT-MG) and isolate genome statistics. Seqs, sequences; avg_cov, average coverage; avg_gc, average G+C content; % rel. abund., percent relative abundance.

**SI Dataset S2.** GCMS measurements of masses 16 (CH_4_), 17 (^13^CH_4_), and 19 (^12^CD_3_H) during the isotope tracing experiment. Percent of labeled methane is calculated as a fraction of provided labeled substrate. Stdev, standard deviation.

**SI Dataset S3.** Gas chromatograph FID measurements of ^12^CH_4_ during isotope tracing experiment. NA, not available/measured.

**SI Dataset S4.** Gas chromatograph FID measurements of CH_4_ during temperature optimum experiment. NA, not available/measured.

**SI Dataset S5.** Inventory of genes expressed by *Ca.* M. hypatiae LCB24 under methanogenic conditions and as depicted in Fig. 5. Expression levels averaged across six replicates are reported in reads per kilobase of transcript per million mapped reads (RPKM).

**SI Dataset S6.** Inventory of genes expressed by Ca. M. hypatiae LCB24 under methanogenic conditions belonging to the beta-oxidation pathway. Expression levels averaged across six replicates are reported in reads per kilobase of transcript per million mapped reads (RPKM).

## References

1. Wolfe JM, Fournier GP. Horizontal gene transfer constrains the timing of methanogen evolution. Nature Ecology & Evolution. 2018;2(5):897–903.

2. Sorokin DY, Makarova KS, Abbas B, Ferrer M, Golyshin PN, Galinski EA, et al. Discovery of extremely halophilic, methyl-reducing euryarchaea provides insights into the evolutionary origin of methanogenesis. Nature Microbiology. 2017;2(8):1–11.

3. Martin WF, Sousa FL. Early microbial evolution: the age of anaerobes. Cold Spring Harbor Perspectives in Biology. 2016;8(2):a018127.

4. Sauterey B, Charnay B, Affholder A, Mazevet S, Ferrière R. Co-evolution of primitive methane-cycling ecosystems and early Earth’s atmosphere and climate. Nature Communications. 2020;11(1):2705.

5. Ueno Y, Yamada K, Yoshida N, Maruyama S, Isozaki Y. Evidence from fluid inclusions for microbial methanogenesis in the early Archaean era. Nature. 2006;440(7083):516-9.

6. Spang A, Ettema TJG. Archaeal evolution: the methanogenic roots of Archaea. Nature Microbiology. 2017;2(8):1–2.

7. Adam PS, Kolyfetis GE, Bornemann TLV, Vorgias CE, Probst AJ. Genomic remnants of ancestral methanogenesis and hydrogenotrophy in Archaea drive anaerobic carbon cycling. Science Advances. 2022;8(44):eabm9651.

8. Wang Y, Wegener G, Williams TA, Xie R, Hou J, Tian C, et al. A methylotrophic origin of methanogenesis and early divergence of anaerobic multicarbon alkane metabolism. Science Advances. 2021;7(27):eabj1453.

9. Hinrichs K-U. Microbial fixation of methane carbon at 2.7 Ga: Was an anaerobic mechanism possible? Geochemistry, Geophysics, Geosystems. 2002;3(7):1–10.

10. Saunois M, Stavert AR, Poulter B, Bousquet P, Canadell JG, Jackson RB, et al. The Global Methane Budget 2000–2017. Earth System Science Data. 2020;12(3):1561–623.

11. Rosentreter JA, Borges AV, Deemer BR, Holgerson MA, Liu S, Song C, et al. Half of global methane emissions come from highly variable aquatic ecosystem sources. Nature Geoscience. 2021;14(4):225–30.

12. Garcia PS, Gribaldo S, Borrel G. Diversity and evolution of methane-related pathways in archaea. Annual Review of Microbiology. 2022;76:727–55.

13. Ferry JG, Kastead KA. Methanogenesis. Archaea: Molecular and Cellular Biology. 2007. p. 288–314.

14. Conrad R. The global methane cycle: recent advances in understanding the microbial processes involved. Environmental Microbiology Reports. 2009;1(5):285–92.

15. Scheller S, Goenrich M, Boecher R, Thauer RK, Jaun B. The key nickel enzyme of methanogenesis catalyses the anaerobic oxidation of methane. Nature. 2010;465(7298):606–8.

16. Thauer RK. Methyl (alkyl)-coenzyme M reductases: nickel F-430-containing enzymes involved in anaerobic methane formation and in anaerobic oxidation of methane or of short chain alkanes. Biochemistry. 2019;58(52):5198–220.

17. Evans PN, Boyd JA, Leu AO, Woodcroft BJ, Parks DH, Hugenholtz P, et al. An evolving view of methane metabolism in the Archaea. Nature Reviews Microbiology. 2019;17(4):219–32.

18. Thauer RK, Kaster AK, Seedorf H, Buckel W, Hedderich R. Methanogenic archaea: ecologically relevant differences in energy conservation. Nature Reviews Microbiology. 2008;6(8):579–91.

19. Baker BJ, De Anda V, Seitz KW, Dombrowski N, Santoro AE, Lloyd KG. Diversity, ecology and evolution of Archaea. Nature Microbiology. 2020;5(7):887–900.

20. Bueno de Mesquita CP, Wu D, Tringe SG. Methyl-based methanogenesis: an ecological and genomic review. Microbiology and Molecular Biology Reviews. 2023;87(1):e00024–22.

21. Söllinger A, Urich T. Methylotrophic methanogens everywhere — physiology and ecology of novel players in global methane cycling. Biochemical Society Transactions. 2019;47(6):1895–907.

22. Nobu MK, Narihiro T, Kuroda K, Mei R, Liu WT. Chasing the elusive Euryarchaeota class WSA2: genomes reveal a uniquely fastidious methyl-reducing methanogen. The ISME Journal. 2016;10(10):2478–87.

23. Vanwonterghem I, Evans PN, Parks DH, Jensen PD, Woodcroft BJ, Hugenholtz P, et al. Methylotrophic methanogenesis discovered in the archaeal phylum Verstraetearchaeota. Nature Microbiology. 2016;1(12):1–9.

24. McKay LJ, Dlakic M, Fields MW, Delmont TO, Eren AM, Jay ZJ, et al. Co-occurring genomic capacity for anaerobic methane and dissimilatory sulfur metabolisms discovered in the Korarchaeota. Nature Microbiology. 2019;4(4):614–22.

25. Wang Y, Wegener G, Hou J, Wang F, Xiao X. Expanding anaerobic alkane metabolism in the domain of Archaea. Nature Microbiology. 2019;4(4):595–602.

26. Evans PN, Parks DH, Chadwick GL, Robbins SJ, Orphan VJ, Golding SD, et al. Methane metabolism in the archaeal phylum Bathyarchaeota revealed by genome-centric metagenomics. Science. 2015;350(6259):434-8.

27. Berghuis BA, Yu FB, Schulz F, Blainey PC, Woyke T, Quake SR. Hydrogenotrophic methanogenesis in archaeal phylum Verstraetearchaeota reveals the shared ancestry of all methanogens. Proceedings of the National Academy of Sciences. 2019;116(11):5037–44.

28. Liu YF, Chen J, Zaramela LS, Wang LY, Mbadinga SM, Hou ZW, et al. Genomic and transcriptomic evidence supports methane metabolism in Archaeoglobi. mSystems. 2020;5(2):e00651–19.

29. Stetter KO, Lauerer G, Thomm M, Neuner A. Isolation of extremely thermophilic sulfate reducers: evidence for a novel branch of archaebacteria. Science. 1987;236(4803):822–4.

30. Slobodkina G, Allioux M, Merkel A, Cambon-Bonavita MA, Alain K, Jebbar M, et al. Physiological and genomic characterization of a hyperthermophilic archaeon *Archaeoglobus neptunius* sp. nov. isolated from a deep-sea hydrothermal vent warrants the reclassification of the genus *Archaeoglobus*. Frontiers in Microbiology. 2021;12:679245.

31. Boyd JA, Jungbluth SP, Leu AO, Evans PN, Woodcroft BJ, Chadwick GL, et al. Divergent methyl-coenzyme M reductase genes in a deep-subseafloor Archaeoglobi. The ISME Journal. 2019;13(5):1269–79.

32. Hua ZS, Wang YL, Evans PN, Qu YN, Goh KM, Rao YZ, et al. Insights into the ecological roles and evolution of methyl-coenzyme M reductase-containing hot spring Archaea. Nature Communications. 2019;10(1):4574.

33. Lynes MM, Krukenberg V, Jay ZJ, Kohtz AJ, Gobrogge CA, Spietz RL, et al. Diversity and function of methyl-coenzyme M reductase-encoding archaea in Yellowstone hot springs revealed by metagenomics and mesocosm experiments. ISME Communications. 2023;3(1):22.

34. Wang J, Qu Y-N, Evans PN, Guo Q, Zhou F, Nie M, et al. Evidence for nontraditional *mcr*-containing archaea contributing to biological methanogenesis in geothermal springs. Science Advances. 2023;9(26):eadg6004.

35. Buessecker S, Chadwick GL, Quan ME, Hedlund BP, Dodsworth JA, Dekas AE. Mcr-dependent methanogenesis in Archaeoglobaceae enriched from a terrestrial hot spring. The ISME Journal. 2023;17(10):1649–59.

36. Laso-Perez R, Krukenberg V, Musat F, Wegener G. Establishing anaerobic hydrocarbon-degrading enrichment cultures of microorganisms under strictly anoxic conditions. Nature Protocols. 2018;13(6):1310–30.

37. Brandis A, Thauer RK. Growth of *Desulfovibrio* species on hydrogen and sulphate as sole energy source. Microbiology. 1981;126(1):249–52.

38. Ai G, Zhu J, Dong X, Sun T. Simultaneous characterization of methane and carbon dioxide produced by cultured methanogens using gas chromatography/isotope ratio mass spectrometry and gas chromatography/mass spectrometry. Rapid Communications in Mass Spectrometry. 2013;27(17):1935–44.

39. Bushnell B. BBMap: a fast, accurate, splice-aware aligner. Lawrence Berkeley National Lab (LBNL), Berkeley, CA (United States); 2014 Mar 17.

40. Seemann T. Prokka: rapid prokaryotic genome annotation. Bioinformatics. 2014;30(14):2068–9.

41. Wu Y-W, Tang Y-H, Tringe SG, Simmons BA, Singer SW. MaxBin: an automated binning method to recover individual genomes from metagenomes using an expectation-maximization algorithm. Microbiome. 2014;2:26.

42. Kang DD, Froula J, Egan R, Wang Z. MetaBAT, an efficient tool for accurately reconstructing single genomes from complex microbial communities. PeerJ. 2015;3:e1165.

43. Alneberg J, Bjarnason BS, de Bruijn I, Schirmer M, Quick J, Ijaz UZ, et al. Binning metagenomic contigs by coverage and composition. Nature Methods. 2014;11(11):1144–6.

44. Miller IJ, Rees ER, Ross J, Miller I, Baxa J, Lopera J, et al. Autometa: automated extraction of microbial genomes from individual shotgun metagenomes. Nucleic Acids Research. 2019;47(10):e57-e.

45. Sieber CMK, Probst AJ, Sharrar A, Thomas BC, Hess M, Tringe SG, et al. Recovery of genomes from metagenomes via a dereplication, aggregation and scoring strategy. Nature Microbiology. 2018;3(7):836–43.

46. Kohtz AJ, Jay ZJ, Lynes MM, Krukenberg V, Hatzenpichler R. Culexarchaeia, a novel archaeal class of anaerobic generalists inhabiting geothermal environments. ISME Communications. 2022;2(1):86.

47. Parks DH, Imelfort M, Skennerton CT, Hugenholtz P, Tyson GW. CheckM: assessing the quality of microbial genomes recovered from isolates, single cells, and metagenomes. Genome Research. 2015;25(7):1043–55.

48. Andrews S, Krueger F, Seconds-Pichon A, Biggins F, Wingett S. FastQC: A quality control tool for high throughput sequence data. Babraham Bioinformatics: Babraham Institute; 2015.

49. Deng Z-L, Münch PC, Mreches R, McHardy AC. Rapid and accurate identification of ribosomal RNA sequences via deep learning. Nucleic Acids Research. 2022;50(10):e60–e.

50. Stahl DA. Development and application of nucleic acid probes in bacterial systematics. Sequencing and hybridization techniques in bacterial systematics. 1991:205–48.

51. Stetter KO. *Archaeoglobus fulgidus* gen. nov., sp. nov.: a new taxon of extremely thermophilic archaebacteria. Systematic and Applied Microbiology. 1988;10:172–3.

52. Huber H, Jannasch H, Rachel R, Fuchs T, Stetter KO. *Archaeoglobus veneficus* sp. nov., a novel facultative chemolithoautotrophic hyperthermophilic sulfite reducer, isolated from abyssal black smokers. Systematic and Applied Microbiology. 1997;20(3):374–80.

53. Mori K, Maruyama A, Urabe T, Suzuki K-i, Hanada S. *Archaeoglobus infectus* sp. nov., a novel thermophilic, chemolithoheterotrophic archaeon isolated from a deep-sea rock collected at Suiyo Seamount, Izu-Bonin Arc, western Pacific Ocean. International Journal of Systematic and Evolutionary Microbiology. 2008;58(4):810–6.

54. Watkins AJ, Roussel EG, Webster G, Parkes RJ, Sass H. Choline and N,N-dimethylethanolamine as direct substrates for methanogens. Applied and Environmental Microbiology. 2012;78(23):8298–303.

55. Hocking WP, Stokke R, Roalkvam I, Steen IH. Identification of key components in the energy metabolism of the hyperthermophilic sulfate-reducing archaeon *Archaeoglobus fulgidus* by transcriptome analyses. Frontiers in Microbiology. 2014;5:95.

56. Li G, Rabe KS, Nielsen J, Engqvist MK. Machine learning applied to predicting microorganism growth temperatures and enzyme catalytic optima. ACS Synthetic Biology. 2019;8(6):1411–20.

57. Mahapatra A, Patel A, Soares JA, Larue RC, Zhang JK, Metcalf WW, et al. Characterization of a *Methanosarcina acetivorans* mutant unable to translate UAG as pyrrolysine. Molecular Microbiology. 2006;59(1):56–66.

58. Ferguson T, Soares JA, Lienard T, Gottschalk G, Krzycki JA. RamA, a protein required for reductive activation of corrinoid-dependent methylamine methyltransferase reactions in methanogenic archaea. Journal of Biological Chemistry. 2009;284(4):2285–95.

59. Estelmann S, Ramos-Vera WH, Gad’on N, Huber H, Berg IA, Fuchs G. Carbon dioxide fixation in ‘*Archaeoglobus lithotrophicus*’: are there multiple autotrophic pathways? FEMS Microbiology Letters. 2011;319(1):65–72.

60. Ferry JG. How to make a living by exhaling methane. Annual Review of Microbiology. 2010;64:453–73.

61. Chernyh NA, Neukirchen S, Frolov EN, Sousa FL, Miroshnichenko ML, Merkel AY, et al. Dissimilatory sulfate reduction in the archaeon ‘*Candidatus* Vulcanisaeta moutnovskia’ sheds light on the evolution of sulfur metabolism. Nature Microbiology. 2020;5(11):1428–38.

62. Neukirchen S, Sousa FL. DiSCo: a sequence-based type-specific predictor of Dsr-dependent dissimilatory sulphur metabolism in microbial data. Microbial Genomics. 2021;7(7).

63. Ramos AR, Keller KL, Wall JD, Pereira IAC. The membrane QmoABC complex interacts directly with the dissimilatory adenosine 5′-phosphosulfate reductase in sulfate reducing bacteria. Frontiers in Microbiology. 2012;3:137.

64. Adams MWW. The metabolism of hydrogen by extremely thermophilic, sulfur-dependent bacteria. FEMS Microbiology Reviews. 1990;6(2-3):219–37.

65. Ma K, Schicho RN, Kelly RM, Adams M. Hydrogenase of the hyperthermophile *Pyrococcus furiosus* is an elemental sulfur reductase or sulfhydrogenase: evidence for a sulfur-reducing hydrogenase ancestor. Proceedings of the National Academy of Sciences. 1993;90(11):5341–4.

66. Hocking WP, Roalkvam I, Magnussen C, Stokke R, Steen IH. Assessment of the carbon monoxide metabolism of the hyperthermophilic sulfate-reducing archaeon *Archaeoglobus fulgidus* VC-16 by comparative transcriptome analyses. Archaea. 2015;2015.

67. Welte C, Deppenmeier U. Membrane-bound electron transport in *Methanosaeta thermophila*. Journal of Bacteriology. 2011;193(11):2868–70.

68. Pereira IA, Haveman SA, Voordouw G. Biochemical, genetic and genomic characterization of anaerobic electron transport pathways in sulphate-reducing delta-proteobacteria. Sulphate-Reducing Bacteria: Environmental and Engineered Systems (LL, B and WA, H, eds), Cambridge University Press, Cambridge, UK. 2007.

69. Grein F, Pereira IA, Dahl C. Biochemical characterization of individual components of the *Allochromatium vinosum* DsrMKJOP transmembrane complex aids understanding of complex function in vivo. Journal of Bacteriology. 2010;192(24):6369–77.

70. Wu K, Zhou L, Tahon G, Liu L, Li J, Zhang J, et al. Isolation of a methyl-reducing methanogen outside the Euryarchaeota. In revision (10.21203/rs.3.rs-2501667/v1).

71. Kohtz A, Krukenberg V, Petrosian N, Jay Z, Pilhofer M, Hatzenpichler R. Cultivation and visualization of a methanogen of the phylum Thermoproteota. In revision (10.21203/rs.3.rs-2500102/v1)

72. Spang A, Caceres EF, Ettema TJG. Genomic exploration of the diversity, ecology, and evolution of the archaeal domain of life. Science. 2017;357(6351):eaaf3883.

## References

1. Buessecker S, Chadwick GL, Quan ME, Hedlund BP, Dodsworth JA, Dekas AE. Mcr-dependent methanogenesis in Archaeoglobaceae enriched from a terrestrial hot spring. The ISME Journal. 2023;17(10):1649–59.

2. Lynes MM, Krukenberg V, Jay ZJ, Kohtz AJ, Gobrogge CA, Spietz RL, et al. Diversity and function of methyl-coenzyme M reductase-encoding archaea in Yellowstone hot springs revealed by metagenomics and mesocosm experiments. ISME Communications. 2023;3(1):22.

3. Apprill A, McNally S, Parsons R, Weber L. Minor revision to V4 region SSU rRNA 806R gene primer greatly increases detection of SAR11 bacterioplankton. Aquatic Microbial Ecology. 2015;75(2):129–37.

4. Bolyen E, Rideout JR, Dillon MR, Bokulich NA, Abnet CC, Al-Ghalith GA, et al. Reproducible, interactive, scalable and extensible microbiome data science using QIIME 2. Nature Biotechnology. 2019;37(8):852–7.

5. Martin M. Cutadapt removes adapter sequences from high-throughput sequencing reads. EMBnetjournal. 2011;17(1):10–2.

6. Callahan BJ, McMurdie PJ, Rosen MJ, Han AW, Johnson AJ, Holmes SP. DADA2: High-resolution sample inference from Illumina amplicon data. Nature Methods. 2016;13(7):581–3.

7. Quast C, Pruesse E, Yilmaz P, Gerken J, Schweer T, Yarza P, et al. The SILVA ribosomal RNA gene database project: improved data processing and web-based tools. Nucleic Acids Research. 2012;41(D1):D590–D6.

8. Davis NM, Proctor DM, Holmes SP, Relman DA, Callahan BJ. Simple statistical identification and removal of contaminant sequences in marker-gene and metagenomics data. Microbiome. 2018;6(1):226.

9. Aramaki T, Blanc-Mathieu R, Endo H, Ohkubo K, Kanehisa M, Goto S, et al. KofamKOALA: KEGG Ortholog assignment based on profile HMM and adaptive score threshold. Bioinformatics. 2020;36(7):2251–2.

10. Lu S, Wang J, Chitsaz F, Derbyshire MK, Geer RC, Gonzales NR, et al. CDD/SPARCLE: the conserved domain database in 2020. Nucleic Acids Research. 2020;48(D1):D265–D8.

11. Sondergaard D, Pedersen CN, Greening C. HydDB: A web tool for hydrogenase classification and analysis. Scientific Reports. 2016;6(1):34212.

12. Blum M, Chang H-Y, Chuguransky S, Grego T, Kandasaamy S, Mitchell A, et al. The InterPro protein families and domains database: 20 years on. Nucleic Acids Research. 2021;49(D1):D344–D54.

13. Neukirchen S, Sousa FL. DiSCo: a sequence-based type-specific predictor of Dsr-dependent dissimilatory sulphur metabolism in microbial data. Microb Genom. 2021;7(7).

14. Neukirchen S, Sousa FL. DiSCo: a sequence-based type-specific predictor of Dsr-dependent dissimilatory sulphur metabolism in microbial data. Microbial Genomics. 2021;7(7).

15. Price MN, Dehal PS, Arkin AP. FastTree 2--approximately maximum-likelihood trees for large alignments. PLoS One. 2010;5(3):e9490.

16. Jay ZJ, Beam JP, Dlakic M, Rusch DB, Kozubal MA, Inskeep WP. Marsarchaeota are an aerobic archaeal lineage abundant in geothermal iron oxide microbial mats. Nature Microbiology. 2018;3(6):732–40.

17. Zaremba-Niedzwiedzka K, Caceres EF, Saw JH, Backstrom D, Juzokaite L, Vancaester E, et al. Asgard archaea illuminate the origin of eukaryotic cellular complexity. Nature. 2017;541(7637):353-8.

18. Edgar RC. MUSCLE: multiple sequence alignment with high accuracy and high throughput. Nucleic Acids Research. 2004;32(5):1792–7.

19. Katoh K, Standley DM. MAFFT multiple sequence alignment software version 7: improvements in performance and usability. Molecular Biology and Evolution. 2013;30(4):772–80.

20. Capella-Gutiérrez S, Silla-Martínez JM, Gabaldón T. trimAl: a tool for automated alignment trimming in large-scale phylogenetic analyses. Bioinformatics. 2009;25(15):1972–3.

21. Minh BQ, Schmidt HA, Chernomor O, Schrempf D, Woodhams MD, von Haeseler A, et al. IQ-TREE 2: new models and efficient methods for phylogenetic inference in the genomic era. Molecular Biology and Evolution. 2020;37(5):1530–4.

22. Rusch A, Amend JP. Order-specific 16S rRNA-targeted oligonucleotide probes for (hyper) thermophilic archaea and bacteria. Extremophiles. 2004;8:357–66.

23. Stahl DA. Development and application of nucleic acid probes in bacterial systematics. Sequencing and hybridization techniques in bacterial systematics. 1991:205–48.

24. Pernthaler A, Pernthaler J, Amann R. Fluorescence in situ hybridization and catalyzed reporter deposition for the identification of marine bacteria. Applied and Environmental Microbiology. 2002;68(6):3094–101.

25. Schaible GA, Kohtz AJ, Cliff J, Hatzenpichler R. Correlative SIP-FISH-Raman-SEM-NanoSIMS links identity, morphology, biochemistry, and physiology of environmental microbes. ISME Communications. 2022;2(1):52.

26. Vaughan RG, Heasler HP, Jaworowski C, Lowenstern JB, Keszthelyi LP. Provisional maps of thermal areas in Yellowstone National Park based on satellite thermal infrared imaging and field observations: US Department of the Interior, US Geological Survey; 2014.

27. Wang Y, Wegener G, Hou J, Wang F, Xiao X. Expanding anaerobic alkane metabolism in the domain of Archaea. Nature Microbiology. 2019;4(4):595–602.

29. Colman DR, Lindsay MR, Amenabar MJ, Boyd ES. The intersection of geology, geochemistry, and microbiology in continental hydrothermal systems. Astrobiology. 2019;19(12):1505–22.

30. Peacock JP, Cole JK, Murugapiran SK, Dodsworth JA, Fisher JC, Moser DP, et al. Pyrosequencing reveals high-temperature cellulolytic microbial consortia in Great Boiling Spring after in situ lignocellulose enrichment. PLoS One. 2013;8(3):e59927.

31. Hua ZS, Wang YL, Evans PN, Qu YN, Goh KM, Rao YZ, et al. Insights into the ecological roles and evolution of methyl-coenzyme M reductase-containing hot spring Archaea. Nature Communications. 2019;10(1):4574.

32. Wang J, Qu Y-N, Evans PN, Guo Q, Zhou F, Nie M, et al. Evidence for nontraditional *mcr*-containing archaea contributing to biological methanogenesis in geothermal springs. Science Advances. 2023;9(26):eadg6004.

33. Klenk H-P, Clayton RA, Tomb J-F, White O, Nelson KE, Ketchum KA, et al. The complete genome sequence of the hyperthermophilic, sulphate-reducing archaeon *Archaeoglobus fulgidus*. Nature. 1997;390(6658):364-70.

34. Slobodkina G, Allioux M, Merkel A, Cambon-Bonavita MA, Alain K, Jebbar M, et al. Physiological and genomic characterization of a hyperthermophilic archaeon *Archaeoglobus neptunius* sp. nov. isolated from a deep-sea hydrothermal vent warrants the reclassification of the genus *Archaeoglobus*. Frontiers in Microbiology. 2021;12:679245.

35. von Jan M, Lapidus A, Glavina Del Rio T, Copeland A, Tice H, Cheng J-F, et al. Complete genome sequence of *Archaeoglobus profundus* type strain (AV18T). Standards in Genomic Sciences. 2010;2(3):327–46.

36. Anderson I, Risso C, Holmes D, Lucas S, Copeland A, Lapidus A, et al. Complete genome sequence of *Ferroglobus placidus* AEDII12DO. Standards in Genomic Sciences. 2011;5(1):50–60.

37. Mardanov AV, Slododkina GB, Slobodkin AI, Beletsky AV, Gavrilov SN, Kublanov IV, et al. The *Geoglobus acetivorans* genome: Fe (III) reduction, acetate utilization, autotrophic growth, and degradation of aromatic compounds in a hyperthermophilic archaeon. Applied and Environmental Microbiology. 2015;81(3):1003–12.

38. Manzella MP, Holmes DE, Rocheleau JM, Chung A, Reguera G, Kashefi K. The complete genome sequence and emendation of the hyperthermophilic, obligate iron-reducing archaeon “*Geoglobus ahangari*” strain 234T. Standards in Genomic Sciences. 2015;10(1):1–19.

39. Stokke R, Hocking WP, Steinsbu BO, Steen IH. Complete genome sequence of the thermophilic and facultatively chemolithoautotrophic sulfate reducer *Archaeoglobus sulfaticallidus* strain PM70-1T. Genome Announcements. 2013;1(4):e00406–13.

40. Boyd JA, Jungbluth SP, Leu AO, Evans PN, Woodcroft BJ, Chadwick GL, et al. Divergent methyl-coenzyme M reductase genes in a deep-subseafloor Archaeoglobi. The ISME Journal. 2019;13(5):1269–79.

41. Mukherjee S, Seshadri R, Varghese NJ, Eloe-Fadrosh EA, Meier-Kolthoff JP, Göker M, et al. 1,003 reference genomes of bacterial and archaeal isolates expand coverage of the tree of life. Nature Biotechnology. 2017;35(7):676–83.

